# Transcriptomic and Lipidomic Profiling of Human iPSC-Derived Microglia Reveals TREM2-Dependent and -Independent Processes

**DOI:** 10.64898/2026.11.16.767071

**Authors:** Christopher L. Cardona, Ronak Patel, Gina M. Finan, Tae-Wan Kim, Andrew A. Sproul, Andrew F. Teich

**Affiliations:** Department of Pathology and Cell Biology, Columbia University Irving Medical Center, New York, NY, USA; Taub Institute for Research on Alzheimer’s Disease and the Aging Brain, Columbia University Irving Medical Center, New York, NY, USA; Department of Neurology, Columbia University Irving Medical Center, New York, NY, USA

**Keywords:** iPSC-microglia, TREM2, transcriptomics, lipidomics, Alzheimer’s disease, amyloid-beta, oligomers, inflammation, cholesterol, DAM

## Abstract

TREM2 has been implicated in AD as a critical regulator of microglial functions and loss of function mutations in TREM2 increases one’s risk of developing AD. Given the diverse functional roles of TREM2, we sought to unravel how it affects downstream pathways using human iPSC-derived microglia-like cells (iMGLs) derived from TREM2 knockout (KO) and isogenic control lines and assessing the response to oligomeric amyloid-β 42 (oAβ42). While previous studies have primarily used fibrillar Aβ, we focused on microglial responses to oAβ42 given their disease relevance and potent cytotoxic effects. At baseline, we identified a set of TREM2-dependent genes, which corresponded to the DAM signature in mice and disease-associated signatures in humans. Additionally, loss of TREM2 resulted in increased expression of cholesterol biosynthesis genes. In response to oAβ42, we found that the transcriptional response was largely TREM2-independent and corresponded to a strong induction of inflammatory pathways, with TREM2 also being downregulated in control iMGLs. We identified subsets of genes and pathways that were unique to either control, such as mitochondrial DNA and RNA degradation, or TREM2 KO iMGLs, such as the senescence-associated secretory pathway and adipogenesis, suggesting some differences in gene expression driven by TREM2 in response to oAβ42. Given that the inflammatory response was TREM2-independent, we sought to identify signaling pathways that contributed to inflammation. Inhibition of toll-like receptor signaling significantly reduced the expression of pro-inflammatory genes and modestly increased expression of TREM2 and TREM2-dependent genes. Consistent with our transcriptional data, cytokine production and secretion in response to oAβ42 was significantly increased to similar levels in both controls and TREM2 KO iMGLs. Lipidomic profiling of microglia revealed that triglycerides and lactosylceramides were significantly increased in both control and TREM2 KO iMGLs in response to oAβ42. TREM2 KO iMGLs had decreased levels of BMPs and increased levels of cholesterol esters across all conditions, further highlighting a role for TREM2 in maintaining lipid homeostasis in microglia. Taken together, we describe genes, pathways, and lipids that are associated with TREM2 expression that likely influence microglial responses to oAβ42.

## 1 INTRODUCTION

Microglia are highly dynamic brain-resident immune cells with diverse functions, such as surveillance, phagocytosis, and immune modulation^1–7^. As the primary phagocytes of the brain, they are responsible for the detection and subsequent removal of apoptotic cells, pathogens, and pathological aggregates, such as amyloid-β (Aβ). Accordingly, microglia have been shown to localize near Aβ plaques in mouse models of Alzheimer’s disease (AD) and human subjects^8–10^, and positively stain for Aβ intracellularly^11^. This highlights that microglia play a central role in responding to Aβ aggregates in AD. Genome wide association studies have repeatedly identified microglia as central mediators of disease in AD^12,13^, with many of these risk genes being primarily expressed in microglia and converging on microglial pathways^14^.

Triggering receptor expressed on myeloid cells 2 (TREM2) is highly expressed in microglia and mediates critical immune functions. TREM2 has been shown to bind Aβ, non-lipidated APOE, phospholipids, and apoptotic debris^15–19^. Following ligand recognition, TREM2 initiates a host of downstream pathways that shape microglial immune functions and responses. Importantly, loss of function mutations in TREM2, specifically the TREM2 R47H mutation, have been associated with an increased risk of developing AD^20^. Single-cell and single-nucleus RNA-sequencing (scRNA-/snRNA-seq) studies in mice and humans have expanded our understanding of how TREM2 mediates the complex microglial responses observed in AD^21–26^. A landmark study using the familial AD mouse model (5xFAD; APP K670N/M671L, APP I716V, APP V717I, PSEN1 M146L, PSEN1 L286V) identified disease-associated microglial (DAM) states and described a trajectory where microglia transition from homeostatic to DAM1 and finally to DAM2 over the course of AD^21^, with the transition from DAM1 to DAM2 being TREM2-dependent, underscoring the importance of TREM2 in regulating this disease-associated response. This study reported that this DAM population was beneficial in AD, representing a microglial subpopulation that emerges in the presence of AD pathology to process and clear plaques^21^. Some human studies have adopted this terminology in human microglial scRNA- and snRNA-seq studies based on genes identified in mice^25^, although it is well-appreciated that there are differences between mouse and human disease-associated signatures^27,28^. Transcriptomic studies in human subjects found that DAM genes were distributed across multiple microglial states rather than one or two distinct states^25,26^. The DAM signature is just one example of how TREM2 regulates microglial responses, and increasing evidence has emerged describing a link between TREM2 and lipid metabolism^29–31^.

As AD progresses, neuronal cell death can lead to accumulation of myelin debris and microglia are required for their clearance^32^. Microglial scavenging of lipid-rich myelin induces gene expression changes related to lipid transport and lipid metabolism that allows for the effective degradation and trafficking of lipids in microglia^33^, including *CD9*, *GPNMB*, *APOE*, *APOC1*, and *LPL*^33,34^. However, chronic accumulation of lipids in microglia impairs their phagocytic capacity^35,36^ and promotes inflammation^36^. For example, one study found that membrane lipids, including glycerophospholipids and sphingolipids, were coregulated during toll-like receptor (TLR) signaling^37^. They found that genetic perturbation of key enzymes involved in sphingolipid metabolism affected TLR-driven IL6 production^37^, highlighting how alterations in membrane lipid composition can affect immune responses. Importantly, TREM2 has been shown to be a critical regulator of cholesterol metabolism during scavenging in mouse models of MS and human induced pluripotent stem cell (hiPSC) microglial models in the context of chronic myelin accumulation^29,30^, emphasizing that TREM2 plays an active role in the regulation of lipid homeostasis in microglia. Furthermore, dysregulated lipid metabolism has been observed in a monocyte-derived microglial model of amyotrophic lateral sclerosis (ALS)^38^, suggesting that lipid-associated changes in microglia may occur across diverse disease contexts. However, whether similar alterations occur in response to Aβ pathology remains poorly understood. Furthermore, the extent to which TREM2 regulates lipid metabolism beyond cholesterol is still unclear, especially under homeostatic conditions.

One of the classical hallmarks of Alzheimer’s disease (AD) is the accumulation of extracellular Aβ. Amyloidogenic processing of amyloid precursor protein leads to the accumulation of Aβ oligomers and fibrils, which ultimately form plaques^39^. Oligomeric forms of Aβ are thought to be neurotoxic and may contribute to downstream disease progression^40^. While oligomeric forms of Aβ are recognized as being neurotoxic in AD, many studies have focused on the microglial responses to fibrillar Aβ^41–44^, emphasizing the importance of better understanding the microglial responses to oligomeric forms of Aβ. Importantly, impaired amyloid clearance has been observed in AD^45^, highlighting this as a key disease process. Moreover, loss of TREM2 in the 5xFAD mouse model showed reduced microglial localization to Aβ plaques and increased amyloid burden^15^.

Given the importance of TREM2, we sought to identify TREM2-dependent functional pathways using transcriptomic and lipidomic approaches. Human induced pluripotent stem cell (hiPSC)-derived microglia-like cells (iMGLs) from TREM2 knockout (KO) and isogenic controls were treated with vehicle or oligomeric Aβ42 (oAβ42) for 6 hours to investigate the acute response to oAβ42. iMGLs mounted a strong pro-inflammatory response in response to oAβ42, which was largely TREM2-independent and instead driven by toll-like receptor (TLR) signaling. In contrast, we identified a set of TREM2-dependent genes including dysregulated cholesterol metabolism genes under steady-state. Lipidomic analyses further confirmed that TREM2 KO microglia had altered cholesterol metabolism in addition to other lipid alterations. These findings support a model in which TREM2 plays a key role in regulating microglial lipid homeostasis and basal functional state, rather than during acute inflammatory responses to oAβ42.

## 2 MATERIALS AND METHODS

### 2.1 Cell culture and microglial differentiations

The FA0000010 (FA10; RUCDR/Infinity BiologiX) and IMR90 cl.4 (WiCell)^46^ hiPSC lines were used in this study under feeder free conditions as described previously^47^. In brief, undifferentiated hiPSCs were maintained in StemFlex media (ThermoFisher) supplemented with penicillin/streptomycin (ThermoFisher) on reduced growth factor Cultrex BME (Biotechne) and routinely split twice a week with ReLeSR (STEMCELL Technologies) without ROCK inhibitor.

To generate microglia, hiPSCs were first differentiated into bankable hematopoietic progenitor cells (HPCs) using the STEMdiff hematopoietic kit (STEMCELL Technologies) largely as per manufacturer’s instructions with minor modifications^48^. HPCs were collected on days 10,12, and 14 and a small sample was taken to validate by flow cytometry using CD43. HPC validation was performed for every HPC differentiation on the first collection. Briefly, cells were washed and immunostained with a CD43-APC antibody (Miltenyi) at 1:100 according to manufacturer’s instructions. Unstained HPCs and negative control hiPSCs were used to set gating for CD43-positive cells on the NovoCyte flow cytometer (Agilent). For cryopreservation, colonies were washed extensively, spun down, and counted before freezing in 45% Medium B, 45% KO serum (ThermoFisher), and 10% DMSO.

Terminal differentiation of HPCs into microglia was done as described previously^49^. HPCs were plated at 200K cells per well on Cultrex-coated 12-well plates in supplemented microglial media. Microglial medium consisted of DMEM/F12 media (ThermoFisher) supplemented with 2X B27 supplement (ThermoFisher), 0.5X N2 supplement (ThermoFisher), 2X insulin-transferrin-selenite ITS-G (ThermoFisher), 1X GlutaMAX (ThermoFisher), 1X MEM non-essential amino acids (ThermoFisher), 1X penicillin-streptomycin (ThermoFisher), 5 µg/mL insulin (Sigma-Aldrich), and 400 µM 1-monothioglycerol (Sigma-Aldrich). For the first 24 days, cells received half-feeds every other day, where fresh media was overlayed without replacement in microglial media supplemented with 100 ng/mL IL-34 (R&D Systems), 50 ng/mL TGFβ1 (STEMCELL Technologies), and 25 ng/mL M-CSF (R&D Systems). On day 12, cells could be split onto a new Cultrex-coated plate, where half of the total cells per well were transferred to a new plate and maintained on the same feeding schedule as the original cells. Cells continued their normal feeding schedule until day 25. On days 25 and 27, microglia received microglial media overlays that was further supplemented with maturation factors, containing 100 ng/mL CX3CL1 (R&D Systems) and 100 ng/mL CD200 (Bon Opus Biosciences), in addition to IL34, TGFβ1, and M-CSF. iMGLs were treated with oAβ42, with or without pretreatment with pharmacological inhibitors, on day 28 for all experiments.

### 2.4 Generation of CRISPRi hiPSCs

IMR90 hiPSCs were treated with dCas9-KRAB-MeCP2 (Addgene; plasmid #122205) lentiviral particles by reverse transduction and selected with 5 µg/mL blasticidin (InvivoGen) to generate permanent lines, using CloneR (STEMCELL Technologies) as described previously^48^. Two independent clones were generated and validated by qPCR for knockdown efficiency using gRNA against the highly expressed cell surface protein EPCAM.

### 2.5 Lentiviral production of TREM2 gRNA

gRNAs targeting TREM2 for CRISPRi was generated by subcloning gRNA sequences into the pLCKO backbone (Addgene; plasmid #73311) as described previously^48^. TREM2 gRNA sequences were kindly provided by the lab of Dr. Alejandro Chavez based on our previous collaborative study^48^. In brief, 1 µg of pLCKO backbone was double digested using BfuAI (NEB) and NsiI (NEB) restriction enzymes to remove the stuffer along with CIP (NEB), which was added post-digestion. Custom DNA oligos (IDT) were used to generate TREM2 gRNA primers targeting 3 different regions in exon 1 of the TREM2 gene with the BfuAI and NsiI restriction sites added to the 5’ and 3’ ends, respectively (**Additional File 9**). Oligos were phosphorylated using T4 PNK (NEB) and annealed in a thermal cycler at 37°C for 30 minutes, 95°C for 5 minutes, and then ramped down to 25°C at 5°C/min. TREM2 gRNA sequences and digested pLCKO were ligated using T4 DNA ligase (NEB) for 1h at RT. 2 µL of reaction was used to transform Stbl3 competent cells (ThermoFisher) and grown on LB-agar plates containing 100 µg/mL ampicillin. Isolated colonies were expanded and confirmed by Sanger sequencing using a U6 primer (Azenta) and then midi-prepped using NucleoBond Xtra Midi Plus kit (Machery-Nagel) for use to generate lentiviral particles.

HEK293T cells were used to generate lentiviral particles, as described previously^47^. 10 million cells were plated onto dishes coated with 0.1% gelatin (Sigma-Aldrich). The next day, cells were transfected using Lipofectamine 3000 (ThermoFisher) with 75 μg of plasmid DNA for 5 hours. Supernatant containing lentiviral particles were collected 48 and 72 hours later. Lentivirus was concentrated using Lenti-X concentrator (Takara Bio) for 1 hour at 4°C. Viral titer was quantified using Lenti-X GoStix plus (Takara Bio).

IMR90 CRISPRi permanent lines differentiated into iMGLs were treated with 5 µL of non-targeting or TREM2 gRNA virus on day 14 of iMGL differentiation. iMGLs received a complete media change 48 hours later and returned to their normal feeding schedule afterwards.

### 2.6 Preparation of amyloid-β (Aβ) oligomers

Recombinant Aβ (1-42) (rPeptide) was used to generate monomers and oligomers, as described previously^49,50^. To generate monomers, Aβ was resuspended in ice cold hexafluoroisopropanol (HFIP; Sigma-Aldrich) to a final concentration of 1 mM using a GasTight syringe (Hamilton) under a fume hood and vortexed for 30 seconds. Aβ solution was aliquoted into protein LoBind tubes (Eppendorf), adding 0.115 mg of Aβ per tube. Aliquots were incubated at room temperature for 2 hours with the lids open to allow for monomerization and evaporation of HFIP. Monomeric Aβ preps were lyophilized by centrifugation with vacuum in a SpeedVac (Sorvall) at 800 xg for 25 minutes at room temperature. Lyophilized product was sealed and stored at −80°C.

For the generation of oligomeric Aβ (oAβ42), aliquots of monomeric Aβ were allowed to come to RT, then resuspended in DMSO to 5 mM of monomeric Aβ. Using a pipette tip, the lyophilized product was thoroughly scraped to ensure complete resuspension and quickly spun down. The sample was sonicated for 10 minutes in a bath sonicator (Brason Ultrasonics) at 18°C. Ice cold phenol-free DMEM/F12 (STEMCELL Technologies) medium was added for a final concentration of 100 µM Aβ. This was incubated for 24 hours at 4°C to promote oligomerization. Prior to treating iMGLs, the oAβ42 was centrifuged at 14,000 xg for 15 minutes at 4°C and the supernatant was used to treat cells at a final concentration of 5µM. Oligomers were tested for endotoxins using the Pierce chromogenic endotoxin quant kit (ThermoFisher).

Aβ oligomers were validated by non-reducing SDS-PAGE. 100-200 ng of oAβ42 was loaded with NuPAGE LDS sample buffer (ThermoFisher) in the absence of reducing agent on a 4-12% NuPAGE bis-tris gradient gel (ThermoFisher) using 1X NuPAGE MES SDS running buffer (ThermoFisher) for 1 hour at 75 V. The was transferred onto 0.2 µm nitrocellulose (GE Healthcare Life Sciences) for 1 hour at 10 V. The membrane was blocked for 1 hour at RT using 5% milk powder (Santa Cruz Biotechnology) diluted in 1X TBS containing 0.1% tween-20 (TBST; Sigma-Aldrich) with gentle rocking. Anti-β-amyloid 6E10 antibody (BioLegend) was diluted 1:1000 in TBST containing 5% BSA (Sigma-Aldrich), added to the membrane, and incubated overnight at 4°C. The following day, the membrane was incubated with 1:5000 HRP-conjugated anti-mouse secondary antibody (ThermoFisher) diluted in TBST containing 5% milk powder for 1 hour at RT. The blot was developed using chemiluminescent HRP substrate and imaged.

### 2.7 Treatment of iMGLs with chemical inhibitors

For toll-like receptor (TLR) inhibition experiments, the inhibitors TAK242 (Tocris) and CU-CPT22 (Tocris) were used to target TLR4 and TLR1/2, respectively. Inhibitors were made up in DMSO and stored at −80°C. iMGLs were pre-treated with 5 μM TAK242 and 20 μM CU-CPT22 for 2 hours prior to treatment with oAβ42.

### 2.8 RNA extraction and qPCR

RNA was extracted using RNeasy Mini Kit (Qiagen) as per manufacturer’s instructions and 100 ng of RNA was used for cDNA synthesis using Maxima first strand cDNA synthesis kit (ThermoFisher). qPCR reactions were prepared using fast SYBR green master mix (ThermoFisher) as per manufacturer’s instructions using target-specific qPCR primers (primer sequences available in **Additional File 9**). Samples were analyzed on a BioRad CFX Opus 384 (BioRad). Fold expression was calculated using the ΔΔC_t_ method and normalized to β-Actin.

### 2.9 Western blotting

Proteins were collected from iMGLs using RIPA buffer (ThermoFisher) supplemented with 1X Halt protease and phosphatase inhibitor (ThermoFisher) as described previously^49^. On ice, cells were scraped and incubated on ice for 10 minutes before centrifuging at 10,000 RPMs for 10 minutes at 4°C. The soluble protein was transferred to a new tube. Protein was quantified using Pierce BCA protein assay kit (ThermoFisher). 8-10 μg of protein was prepared in NuPAGE LDS buffer (ThermoFisher) and NuPAGE denaturing buffer (ThermoFisher) and boiled for 10 minutes at 70°C. Samples were run on a 4-12% NuPAGE bis-tris gradient gel (ThermoFisher) using 1X NuPAGE MES SDS running buffer or 1X NuPAGE MOPS SDS running buffer (ThermoFisher) and transferred onto 0.2 μm or 0.45 μm nitrocellulose membranes. Western blotting was performed similarly to that described in the section for preparing amyloid-β oligomers. Anti-TREM2 antibody (Cell Signaling Technologies) was used at 1:1000 and 1:5000 HRP-conjugated anti-rabbit secondary antibody was used. β-Actin (Cell Signaling Technologies) or GAPDH (Cell Signaling Technologies) were used as loading controls at 1:1000.

### 2.10 Immunofluorescence

iMGLs were grown on Cultrex-coated coverslips for immunofluorescence staining as described previously^49^. Cells were gently washed two times with PBS before fixing with 4% PFA (ThermoFisher) for 15 minutes at RT. Cells were washed three times with PBS. Blocking was performed using SuperBlock PBS (ThermoFisher) supplemented with 0.1% Triton X-100 (Sigma-Aldrich) for 1 hour at RT. Anti-IBA1 (Wako) and anti-P2RY12 antibodies were used at 1:250 and 1:300, respectively, and incubated at 4°C overnight. The following day, cells were washed three times before adding Alexa Fluor-conjugated rabbit anti-mouse secondary antibodies at 1:1000 for 1 hr at RT protected from light. Cells were washed three times, stained with DAPI (ThermoFisher) for 10 mins, and mounted using Vectashield hard mount antifade mounting medium (Vector Laboratories). Images were taken using the ECHO revolve 2 microscope (Discovery Echo).

### 2.11 Cytokine quantification

Following treatment of iMGLs with oAβ42, conditioned media (CM) was collected from individual wells into 1.5 mL Eppendorf tubes. CM was centrifuged at 1200 RPMs for 3 min. at 4°C and supernatant was aliquoted at 200 μL each. CM was stored at −80°C. Total protein in the CM was quantified using the 660 nm protein assay (ThermoFisher) as per manufacturer’s instructions. 100 μL of CM was sent to Eve Technologies and 15 different analytes were quantified using the human cytokine proinflammatory focused 15-plex discovery assay (Eve Technologies) in duplicate for every sample. Raw data were normalized to total protein in the CM and reported in pg/mL.

### 2.12 Bulk RNA-seq analyses

RNA was collected from iMGLs on day 28 post HPC following treatment. For bulk RNA-sequencing (RNA-seq), 150-300 ng of RNA was submitted for paired-end RNA-seq at 30 million read depth (Azenta Life Sciences). Three experimental replicates (with 3 technical replicates each) were sent for sequencing separately for FA10 TREM2 KO iMGLs and one experimental replicate (with 3 technical replicates) was sent for sequencing for IMR90 CRISPRi iMGLs. Raw FASTQ files were aligned to the human transcriptome (Ensembl assembly version 113) using STAR (version 2.7.10b) and standard ENCODE parameters. Counts were generated using featureCounts from subread (version 2.0.1) on the aligned bam files. Raw counts were imported into R (version 4.3.2) and all downstream analyses were performed in R.

#### Data pre-processing and quality control

Data pre-processing was performed to remove genes with low counts and remove outliers. Genes with less than 10 reads in 15% of the samples were removed from the analysis. Individual replicate variability was assessed using MA-based sample comparisons on variance-stabilized expression data and implemented with RNAseqQC (version 0.2.1)^51^. MA plots were used to visualize fold change differences (M values) and average gene expression (A values) between a sample and its group reference. Sample-level transcriptional deviations were quantified using the median absolute deviation (MAD) of gene-wise expression differences between an individual sample and the corresponding experimental group, where samples with MAD values above 0.5 were excluded. Only one sample out of 36 was removed from the downstream analysis due to high variability (**Additional File 2A-B**). Processed raw data was used for differential expression analysis. Principal component analysis (PCA) and sample clustering were used to assess experimental sources of variation across experiments. Variance stabilized and batch corrected data were used for PCA. Raw data were normalized using the variance stabilizing transformation^52^ through DESeq2 (version 1.42.1)^53^. Further, batch effects arising from independent sequencing runs were regressed using removeBatchEffect from limma (3.58.1). Ensembl gene IDs were converted to gene symbols using biomaRt (version 2.58.2) and org.Hs.eg.db (version 3.18.0) packages with only protein-coding genes being carried forward for downstream analysis.

#### Differential gene expression

Differential expressed genes (DEGs) were identified using DESeq2 (version 1.42.1)^53^, where the processed raw counts, metadata, and design matrix were converted into a DESeq object. The design matrix was generated to account for batch effects, TREM2 effects, treatment effects, and the interaction of TREM2 and treatment effects (TREM2:treatment). To run DESeq2, estimateSizeFactors was used, then DESeq. Contrasts were specified to derive DEGs for each comparison of interest. Significant DEGs were those with a false discovery rate (FDR) less than 0.05. Volcano plots were generated using EnhancedVolcano (version 1.20.0) with standard parameters. For visualization of gene expression across all samples, heatmaps were generated using pheatmap (version 1.0.13). Variance stabilized and batch corrected data were used to generate z-scores and subsequent visualization via heatmap. For the visualization of overlapping and unique DEGs between isogenic control and TREM2 KO iMGLs in response to oAβ42, ggvenn (version 0.1.10) was used with only significant DEGs being compared, those with a FDR less than 0.05.

#### Pathway analysis

For analysis of TREM2-related changes under steady-state conditions, fGSA (version 1.28.0)^54^ was used to perform pathway analysis. Differential gene expression changes were ranked according to the DESeq2 Wald test statistic. Pathways from Reactome 2025^55^ were used to perform pathway analysis. The top 10 enriched pathways for both isogenic control or non-targeting control and TREM2 KO or TREM2 knockdown (KD) iMGLs were selected based on normalized enrichment score. Top pathways were visualized using ggplot2 (version 4.0.0).

For analysis of gene expression changes in response to oAβ42, enrichR (version 3.4)^56^ was performed using DEGs from the isogenic or non-targeting control response and TREM2 KO or TREM2 KD response to oAβ42. Only significant DEGs were used for pathway analysis, those with a FDR less than 0.05. These DEGs were used as input for enrichR^56^ pathway analysis using the gene ontology biological processes database. Plots were generated using ggplot2. For isogenic control- and TREM2 KO-specific DEGs, the setdiff base R command was used. These were used as input for enrichR^56^ to perform pathway analysis.

#### Log fold change analysis

Log_2_ fold changes from DESeq2 for isogenic control and TREM2 KO iMGLs in response to oAβ42 were combined to compare gene expression changes. A distance metric was utilized to quantify how different the log_2_ fold expression changes were between groups. To calculate the distance, the fold change in isogenic control iMGLs was subtracted from the fold change in TREM2 KO iMGLs to provide an absolute value for the fold change difference. The absolute fold change difference was divided by the square root of two to generate a distance metric for every gene. Those with an absolute distance of at least 0.7, which corresponds to a minimum log_2_ fold change of one, were selected for further analysis. Genes that passed this threshold were used as input for pathway analysis using enrichR^56^.

#### Gene set score analysis

Marker genes were extracted from mouse DAM signatures^21^, human AD microglial signatures^24^, human DAM subclusters^25^, and xenotransplanted iMGLs^57^. For human signatures and xenotransplanted iMGLs, the top significant cluster-defining genes were extracted across all clusters and used to define the corresponding microglial state, setting a consistent number of genes to use based on the lowest number of genes that defined any given cluster. A gene set score was calculated using the normalized and scaled data. This was defined as the average z-score per sample for all genes in the gene set. Gene set scores were plotted as box plots showing all points with the error bars representing the minimum and maximum values.

### 2.13 Lipidomic analyses

On day 28, after oAβ42 treatment, media was collected and centrifuged to collect cells in suspension. For attached cells, 500 μL of Accutase was added to dissociate cells. After resuspending in base microglial media, the cell pellets were combined to a total volume of 1 mL. Live cells were counted using the Countess II automated cell counter (ThermoFisher). 200,000 to 250,000 cells (same number of cells were used for all replicates within an experiment) were aliquoted into a 1.5 mL microcentrifuge tube and centrifuged at 4°C. Cell pellets were stored at −80°C before submitting to the Columbia University Biomarkers Core Laboratory for analysis of over 590 individual lipid species. Lipids were extracted from cell pellets using a modified Bligh-Dyer method after supplementation with internal standards. Separation of extracted glycerophospholipids and sphingolipids was performed using normal phase high-performance liquid chromatography (HPLC), while sterols and glycerolipids were separated on a reverse-phase column. Lipidomics was performed using the 6490 Triple Quadrupole mass spectrometer integrated with an Agilent 1260 Infinity liquid chromatography system. Lipid species were quantified using dynamic multiple reaction monitoring. During sample processing, a small aliquot was used for total protein quantification using the BCA protein assay as per manufacturer’s instructions. Total protein was used to normalize lipidomic results. The molar percentages and normalized lipid concentrations were used for data analysis performed in R (version 4.3.2). Lipid species whose abundance was zero across all samples were removed from downstream analyses. Principle component analysis (PCA) was used to assess the contribution of genotype and treatment to overall variance in lipid abundance and was visualized using lipidr (version 2.16.0)^58^.

#### Lipid compositional analysis

Normalized lipid concentrations were scaled and averaged per experimental group, then visualized by heatmap using Prism (version 10.0.2, GraphPad). Lipids were stratified by broad lipid classes, including fatty acyls, glycerophospholipids, sphingolipids, and sterol lipids based on the LIPID MAPS classification system^59,60^ and molar percentages were averaged per experimental group to characterize the lipid composition across samples.

#### Differential lipid abundance

Raw molar percentages were normalized by log transformation. A linear model was fit to each lipid using lmFit in limma (version 3.58.1), with a design matrix accounting for experimental factors. Empirical Bayes moderation was applied to the linear model fit using eBayes in limma to shrink lipid-wise residual variances. Contrasts were designated to yield differential lipid abundances. The log_2_ fold changes were then compared between genotypes and plotted using ggplot2 (version 4.0.0) to assess differences in lipid abundances in response to oAβ42. Specific lipid species were highlighted that had a log_2_ fold change difference of at least one. To visualize lipid species abundances for a specific lipid class across samples, the log transformed abundances were scaled, then plotted on a heatmap using pheatmap (version 1.0.13).

### 2.14 Statistical Analyses

Some data were analyzed using GraphPad prism. The appropriate test was used as per the experimental design: unpaired t-test when comparing two groups and two-way ANOVA when studying the effect of two independent variables. For unpaired t tests, a Shapiro-Wilk test was used to assess normality, and an F test was used to compare variances. If the variance was different between groups, an unpaired t test with Welch’s correction was used. If the data were nonparametric, the Mann-Whitney test was used in place of the t-test. For two-way ANOVA, a full model was used to identify interaction effects. Following a significant ANOVA, Tukey’s post hoc tests were performed to identify specific pairwise differences between groups at a 95% confidence level.

## 3 RESULTS

### 3.1 Loss of TREM2 induces cholesterol biosynthesis-related genes and disrupts the DAM transcriptional signature

To study human microglia, we used a previously established protocol for generating hiPSC-derived microglia-like cells (iMGLs)^61^, as we have done previously^48,49^. iMGLs generated from FA0000010 (FA10) hiPSCs^48,49^ exhibited low expression of hiPSC-markers (**Additional File 1A-B**), increased expression of key microglial markers (**Additional File 1C-F**), high surface expression of CD45 and CD11b (**Additional File 1G**), and expressed IBA1 and P2RY12 as measured by immunofluorescence (**Additional File 1H-I**). We utilized FA10 TREM2 KO and isogenic control (CTRL) hiPSCs, which were kindly provided by Dr. Tae Wan-Kim^62^, and differentiated them into iMGLs to investigate how TREM2 influences microglial functions. We confirmed the loss of TREM2 at the protein level in iMGLs via western blot (**Additional File 1K**). To better understand the role of TREM2 in regulating microglial processes, we conducted 3 independent differentiations of TREM2 KO and isogenic control hiPSCs into iMGLs, treated with either vehicle (F12) or 5µM oAβ42 and were subjected to bulk RNA-sequencing (RNA-seq).

We first assessed the differences between TREM2 KO and CTRL iMGLs under baseline conditions (vehicle). We observed 703 differentially expressed genes (DEGs) between KO and CTRLs, with a majority being downregulated (**Figure 1A**; see full list of DEGs in **Additional File 3**). Many of the genes downregulated in TREM2 KO iMGLs were canonical DAM genes, including *MITF, GPNMB, CD9, APOE,* and *LPL*, suggesting that these are TREM2-dependent genes (**Figure 1A**). This aligns well with original observations in mice where the transition from homeostatic microglia to DAM (at least the DAM2 state) was TREM2-dependent^21^. Additionally, we detected cholesterol metabolism-related genes being upregulated in KO iMGLs, such as *SREBF2, HMGCS1,* and *SQLE* (**Figure 1A**). Next, we performed pathway analysis, which showed that the most enriched pathway in KO iMGLs was cholesterol biosynthesis (**Figure 1B**; see full pathway analysis results in **Additional File 3**). Furthermore, regulation of cholesterol biosynthesis by *SREBF2* was also enriched in KO iMGLs (**Figure 1B**), suggesting a relationship between TREM2 and cholesterol metabolism at baseline. This increase in cholesterol biosynthesis genes was accompanied by downregulation of cholesterol efflux genes (**Figure 1C**), highlighting dysregulated cholesterol processing in KO iMGLs. We also observed an enrichment of pathways related to DNA damage responses and proliferation (**Figure 1B**), supporting a previously identified role for TREM2 signaling pathways in cell cycle regulation^63,64^. Pathways that were downregulated in TREM2 KO iMGLs included interferon signaling, insulin receptor signaling, and MITF-driven regulation of genes related to lysosome biogenesis and autophagy (**Figure 1B**). This is consistent with the increased expression of DAM genes, as they have been functionally related to lysosomal function previously^65,66^. We validated these results using an independent system for studying reduced TREM2 function. Using CRISPR inhibition (CRISPRi) via a permanent line in IMR90 hiPSCs that expressed dCas9-KRAB-MeCP2 constitutively^67^, we treated iMGLs with guide RNAs (gRNAs) targeting TREM2 or non-targeting (NT) control via lentiviral transduction, resulting in decreased TREM2 gene expression. Pathway analysis using this model revealed similar changes between NT and TREM2 knockdown (KD) iMGLs as was seen in TREM2 KO versus its isogenic control (see full list of DEGs and pathway analysis results in **Additional File 3**). Specifically, we saw an enrichment of pathways and genes relating to cholesterol biosynthesis (**Additional File 4A**). We also similarly observed a downregulation of cholesterol efflux genes in TREM2 KD iMGLs (**Additional File 4B**), further supporting a role for TREM2 in regulating cholesterol homeostasis. Overall, these analyses revealed an important set of TREM2-dependent genes and that loss of or reduced TREM2 leads to altered expression of cholesterol-related genes and pathways.

**Figure 1:**
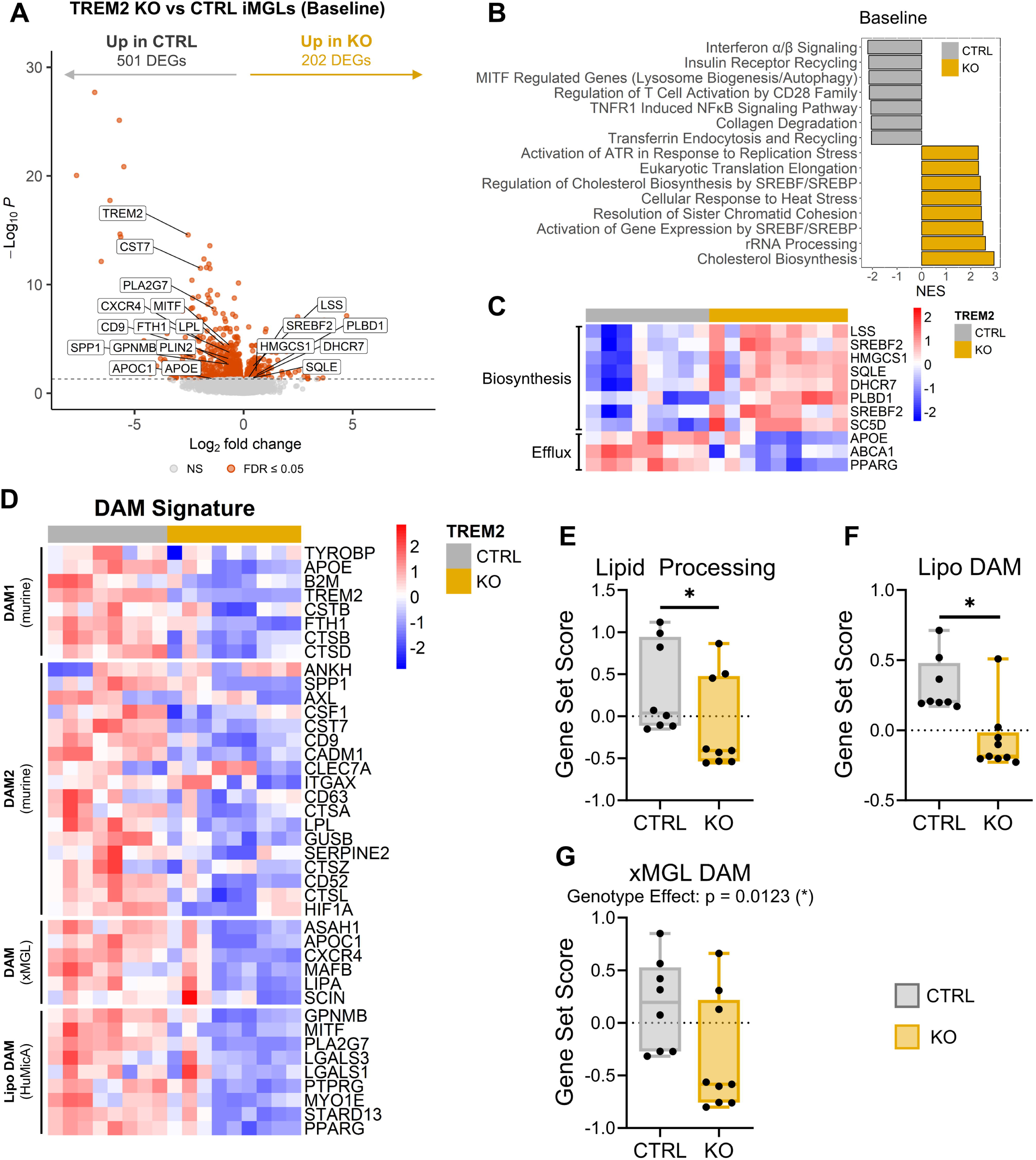
Transcriptional Analysis of TREM2 KO iMGLs at Steady-State. (**A**) Volcano plot comparing TREM2 KO and CTRL iMGLs at baseline, with orange points representing significant DEGs, which pass the threshold for significance (FDR ≤ 0.05). DEGs with a negative log_2_ fold change were more highly expressed in CTRL iMGLs and those with a positive log_2_ fold change were more highly expressed in TREM2 KO iMGLs. (**B**) Pathway analysis results using gene set enrichment analysis, showing the top pathways ranked by normalized enrichment score (NES), where a positive NES corresponds to enriched pathways in KO (yellow) and negative NES corresponds to enriched pathways in CTRLs (grey) using Reactome. (**C**) Heatmap of normalized z-scores for genes related to cholesterol biosynthesis and efflux. (**D**) Heatmap showing the normalized z-score gene expression in CTRL and TREM2 KO iMGLs of DAM genes described in mice (Keren-Shaul *et al*.), xenotransplanted microglia (xMGL; Mancuso *et al*.), and human microglia (HuMicA; Martins-Ferreira *et al*.). (**E-G**) Gene set score analysis using the top 40 cluster-defining genes from xMGL DAM cluster (**E**), lipid-associated DAM cluster from human microglia (**F**; Lipo DAM; Martins-Ferreira *et al*.), and lipid-processing cluster from human microglia in AD (**G**; Sun *et al*.). Significance testing was performed using two-way ANOVA on the full dataset, with significant results from two-way ANOVA being shown if the TREM2 effect was significant (* p-value ≤ 0.05, ** p-value ≤ 0.01, *** p-value ≤ 0.001). Boxplots show data from the 25th to the 75th percentile, with the whiskers extending to the minimum and maximum values and the line in the box corresponding to the median. Sequencing data was obtained from 3 independent experiments.

Next, we sought to validate these TREM2-dependent genes by cross-comparing with previously published single-cell RNA-sequencing datasets which described DAM clusters. First, we wanted to compare our findings with those originally observed by Keren-Shaul *et al* in AD (5X FAD) mice^21^. They described a transition of murine AD microglia from homeostatic to DAM1 to DAM2, with the transition from DAM1 to DAM2 being TREM2-dependent. We observed a downregulation of both DAM1 and DAM2 genes in our TREM2 KO iMGLs (**Figure 1D**), suggesting that both sets of DAM genes may be TREM2-dependent at baseline in human microglia. Note that while the initial description of murine DAM genes was under pathological conditions, this data examines these same genes in human CTRL iMGLs under basal conditions, in addition to oAβ42 treatment conditions. Importantly, others have previously shown that DAM genes are downregulated in TREM2 KO iMGLs compared to controls under basal conditions, such as *APOE*, CD9, *GPNMB*, *PLA2G7*, *APOC1*, and *LPL*^30,68^. This is further supported by our previous study in iMGLs at the single-cell level, where we observed stable expression of the iMGL DAM cluster under untreated conditions^48^. We then wanted to know how closely our TREM2-dependent genes mirrored human disease-associated signatures. We used three independent datasets: human scRNA- and snRNA-seq of microglia across neurological conditions (including AD, autism spectrum disorder, multiple sclerosis, epilepsy, Lewy body disease, COVID-19, and healthy controls - HuMicA)^25^, human snRNA-seq of microglia in AD across 400 human patient samples and controls^24^ (Sun *et al*.), and human iMGLs xenotransplanted into the *APP^NL-G-F^* AD mouse model (xMGLs)^57^. We found that human disease-associated signatures were expressed at lower levels in our TREM2 KO iMGLs (**Figure 1D**). Additionally, in our TREM2 KD model, we observed a similar downregulation of DAM genes described in mouse and human studies^21,25,57^ (**Additional File 4C**), supporting a role for TREM2 in regulating disease-associated genes even under basal conditions. snRNA-seq from human microglia (Sun *et al*.) identified a lipid processing cluster that closely resembled the classical DAM signature described in mice^24^, which we show as being significantly downregulated in our data (**Figure 1E**). Microglia from the HuMicA study^25^ were described as three functionally distinct human disease-associated populations, lipid-associated DAM (Lipo DAM), inflammatory DAM, and ribosomal-associated DAM. Using gene set score analysis, we observed a significant downregulation of only the Lipo DAM gene signature (**Figure 1F**). For xMGLs^57^, we observed a genotype-specific effect with decreased expression of the DAM signature in our KO iMGLs (**Figure 1G**).

Taken together, we describe a set of TREM2-dependent genes which largely overlap with murine DAM genes as well as human disease-associated gene sets. Intriguingly, these changes in gene expression are observed even under basal conditions, suggesting that these genes are not only involved in disease, but in normal microglial function. Importantly, we observed dysregulation of transcriptional programs related to cholesterol metabolism, suggesting a role for TREM2 in balancing cholesterol biosynthesis and efflux. While TREM2 has previously been implicated in cholesterol metabolism during scavenging of myelin debris, this work was primarily done in a mouse model of demyelination^29^. The findings presented here highlight that TREM2 may play a functional role in regulating cholesterol in microglia, even in the absence of exogenous alterations in lipid abundance.

### 3.2 Microglial pro-inflammatory responses to oligomeric amyloid-β are largely TREM2-independent

Next, we wanted to understand how TREM2 was involved in shaping the response to oligomeric amyloid-β 1-42 (oAβ42). We treated iMGLs with either vehicle or oAβ42 for 6 hours and subjected cells to RNA-seq. oAβ42 preparations were validated before every experiment to confirm the generation of oligomers (**Additional File 1J**). Principal component analysis (PCA) revealed that the strongest source of transcriptomic variation was related to oAβ42 treatment (91% of variance), with no clear separation observed based on TREM2 genotype (**Additional File 2C**). We identified many DEGs in response to oAβ42, which included 5,538 upregulated genes and 5,660 downregulated genes (**Figure 2A-B**). Interestingly, we found that many of these changes in gene expression were largely similar between CTRL and TREM2 KO iMGLs (**Figure 2A-B**; see full list of DEGs in **Additional File 3**), consistent with our PCA results (**Additional File 2C**). We found similar changes in gene expression in response to oAβ42 in our CRISPRi TREM2 KD model (**Additional File 4D-E**), with many of the upregulated and downregulated genes being shared between TREM2 KD iMGLs and non-targeting controls. Many of the upregulated pathways were related to inflammatory activation, including cytokine signaling, signaling by interleukins, antigen presentation, and activation of NFκB (**Figure 2C**; see full pathway analysis results in **Additional File 3**), suggesting that human microglia mount an early strong inflammatory response against amyloid oligomers. Downregulated pathways included those associated with GTPase signaling, lipid metabolism, and DNA repair (**Figure 2D**). This decrease in GTPase signaling is likely reflective of microglia losing their homeostatic functions, as GTPase signaling has been associated with homeostatic microglia^25^. We verified that genes associated with cytokine signaling and lipid metabolism pathways were changing in response to oAβ42 (**Figure 2E-F**).

**Figure 2:**
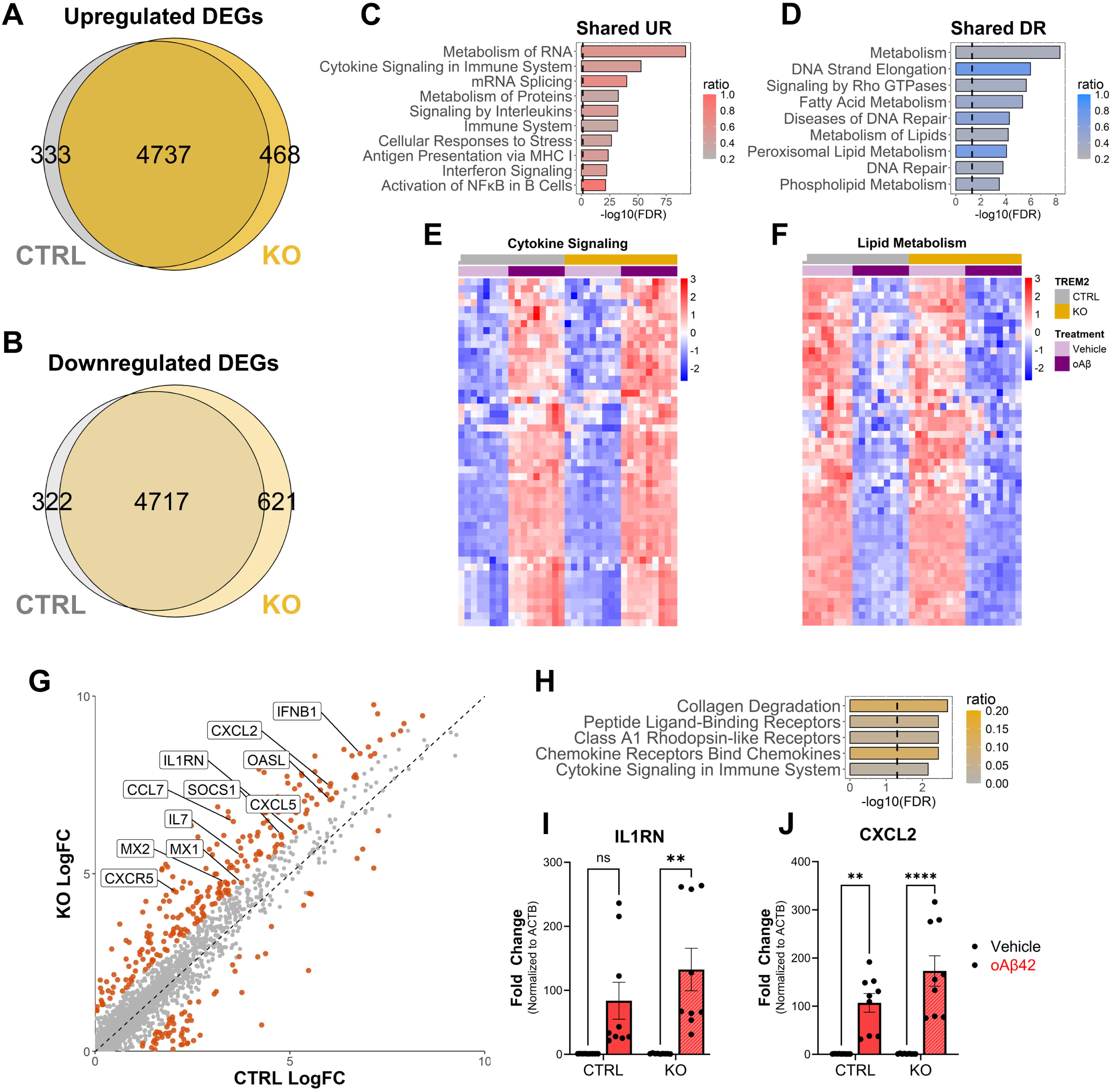
Transcriptomic Responses to Oligomeric Aβ42 in CTRL and TREM2 KO iMGLs. (**A**) Venn diagram comparing DEGs that were upregulated (**A**) and downregulated (**B**) in response to oAβ42 in CTRL (grey) and TREM2 KO (yellow) iMGLs. Only significant genes are shown with no threshold for log_2_ fold change. (**C-D**) Pathway analysis using Reactome terms for shared upregulated (**C**) and downregulated pathways (**D**), with the dotted line representing minimum significance of FDR ≤ 0.05 and the color of the bar representing the ratio of the number of input genes divided by the total number of genes in that pathway. (**E-F**) Heatmaps show the normalized z-score expression of genes belonging to that pathway. (**G**) For every gene, the log_2_ fold change in response to oAβ42 was plotted with the fold change in CTRL on the x-axis and for TREM2 KO on the y-axis. Points highlighted in orange were genes that had at least a two-fold difference in gene expression when comparing to either CTRL or KO iMGLs. (**H**) Those genes with at least a two-fold higher expression in response to oAβ42 in TREM2 KO iMGLs (orange points above the dotted line) was used for pathway analysis using Reactome pathways, with the dotted line representing minimum significance of FDR ≤ 0.05 and the color of the bar representing the ratio of the number of input genes divided by the total number of genes in that pathway. (**I-J**) mRNA expression of *IL1RN* (**C**) and *CXCL2* (**D**) validated by qPCR. Two-way ANOVA was used for statistical significance testing (** p-value ≤ 0.01, **** p-value ≤ 0.0001), followed by Tukey’s post hoc analysis. Error bars are shown as SEM.

While we observed largely overlapping changes in gene expression in response to oAβ42, we wanted to identify shared upregulated genes that may be enhanced by loss of TREM2. To this end, we calculated a distance metric for every gene when comparing the log_2_ fold change in CTRL and KO iMGLs and identified 291 upregulated genes with differential induction (**Figure 2G**). We highlighted 230 genes that had at least a two-fold higher expression in KO iMGLs in comparison to CTRL iMGLs (**Figure 2G**) and found that these genes were associated with collagen degradation, chemokine receptors, and cytokine signaling (**Figure 2H**). We validated these findings via qPCR and found that *IL1RN* and *CXCL2* had higher expression in KO iMGLs (**Figure 2I-J**), suggesting that while there was a strong upregulation of pro-inflammatory genes in both CTRL and KO iMGLs, TREM2 may play a more nuanced role in regulating immune genes in response to oAβ42.

We used gene set score analysis and compared our data with human microglial data from AD patients and controls to contextualize how our transcriptomic responses to oAβ42 relate to human signatures. Sun *et al*. characterized one of the largest human microglial datasets from patients with AD by snRNA-seq^24^. We found that homeostatic (MG0) gene expression was downregulated in response to oAβ42 similarly between CTRL and KO iMGLs (**Additional File 5A**). Importantly, we observed a significant reduction in genes related to neuronal surveillance (MG1; **Additional File 5B**), which was significantly downregulated in human subjects with AD, suggesting a vulnerability in human microglia that may be exacerbated by loss of TREM2. Sun *et al*. reported three inflammatory clusters, which corresponded to different levels of inflammatory activation which were functionally categorized based on inflammatory pathway enrichment, with cluster III (MG10) having the strongest enrichment and cluster II (MG8) having the lowest enrichment^24^. Inflammatory clusters I and III were highly upregulated in response to oAβ42 in both CTRL and KO iMGLs (**Additional File 5C, J**), confirming that microglia initiate a strong pro-inflammatory response acutely, which is significantly higher in KO iMGLs. This highlights that TREM2 may play a role in fine-tuning the expression of pro-inflammatory genes in response to pathology. The weakest inflammatory cluster II was downregulated in response to oAβ42 in both CTRL and KO iMGLs (**Additional File 5I**). Additionally, we observed an upregulation of glycolytic genes in response to oAβ42 in CTRL and KO iMGLs, which may reflect metabolic changes that often accompany immune activation (**Additional File 5H**). TREM2 KO iMGLs had significantly lower expression of glycolytic genes at baseline compared to controls, suggesting that loss of TREM2 may alter basal metabolic states in iMGLs (**Additional File 5H**). Lastly, we observed a significant decrease in genes related to phagocytosis, but only in KO iMGLs (**Additional File 5F**), which is in line with in vitro and in vivo studies showing that TREM2 deletion or suppression reduces microglial phagocytosis^62,68–70^.

We then asked whether DAM genes were affected in TREM2 KO iMGLs in response to oAβ42. We used human microglial disease-associated signatures across three studies: MG4 lipid processing cluster (**Additional File 5E**; Sun *et al*.), DAM subclusters from the human microglial atlas (**Additional File 5M-O**), and the DAM cluster from xMGLs (**Additional File 5P**). We observed a treatment-specific increase in the inflammatory DAM gene signature (**Additional File 5M**), which was similarly increased in both CTRL and KO iMGLs. However, we saw no significant changes in lipid associated DAM, ribosome-associated DAM, or DAM from xMGLs in response to oAβ42 (**Additional File 5N-P**). Although these additional DAM clusters have been tied to AD pathology, our data suggest this is not the case for acute stimulation with oAβ42 in our model system.

Overall, we show that there are large transcriptional changes during the acute response to oAβ42, with the majority of them being TREM2-independent in our model system. In particular, induction of the pro-inflammatory response does not require TREM2. However, rather than acting as a driver of inflammation, TREM2 seems to subtly alter the expression of specific immune genes, such as *IL1RN* and *CXCL2*, which may lead to differences in immune activation. When comparing our data with human AD microglial inflammatory signatures, we also observe significantly higher expression levels of inflammatory clusters I (MG2) and III (MG10) in TREM2 KO iMGLs compared to controls. Additionally, changes in glycolytic metabolism and phagocytosis were observed and seem to be differentially affected by loss of TREM2, highlighting that TREM2 still participates to some degree in diverse transcriptional responses to oAβ42 in microglia relating to immune signaling, metabolism, and phagocytosis.

### 3.3 Characterizing control- and TREM2 KO-specific responses to oligomeric amyloid-β

While there were many similar transcriptional changes in response to oAβ42 between CTRL and TREM2 KO iMGLs, we wanted to know what biological processes were associated with genes that were specifically upregulated in either CTRL or TREM2 KO iMGLs in response to oAβ42. We observed 333 CTRL-specific and 468 KO-specific upregulated DEGs (**Figure 2A**). For CTRL-specific responses, we found that many pathways were associated with mitochondrial RNA processing and these genes were uniquely upregulated in CTRL iMGLs in response to oAβ42, but not KO iMGLs (**Additional File 6A-B**). During inflammatory processes, such as those mediated by oAβ42, mitochondria can experience cellular stress leading to mitochondrial DNA or RNA release, which can be detected by multiple intracellular proteins to promote aberrant pro-inflammatory processes^71,72^. KO-specific upregulated DEGs were associated with pathways related to adipogenesis, viral RNA regulation, and senescence-associated secretory phenotype (SASP), with these genes being specifically upregulated only in TREM2 KO iMGLs in response to oAβ42 (**Additional File 6C-D**). Increased expression of genes relating to adipogenesis in KO iMGLs, such as *CREBBP* and *PLIN4*, may reflect alterations in lipid storage within microglia. As cholesterol metabolism seems to be dysregulated at baseline in TREM2 KO iMGLs, exposure to oAβ42 may drive lipid accumulation in microglia lacking TREM2.

Overall, we identified specific microglial processes that were affected by loss of TREM2. Importantly, this included upregulation of genes associated with adipogenesis in KO iMGLs. Additionally, CTRL iMGLs exhibited an upregulation of mitochondrial DNA regulation genes, highlighting how inflammation may drive mitochondrial stress, which needs to be tightly regulated in microglia to avoid activation of additional inflammatory pathways. These data provide evidence that loss of TREM2 in iMGLs results in a distinct transcriptomic response to oAβ42 compared to controls, which involves changes in gene expression related to adipogenesis and cellular senescence.

### 3.4 Toll-like receptor signaling drives inflammatory responses to oligomeric amyloid-β

We observed many significant shared upregulated pathways (679) between CTRL and KO iMGLs in response to oAβ42, many of which were associated with pro-inflammatory signaling (**Additional File 3**). Some of these included TLR signaling and NFκB signaling pathways, which are part of the classical TLR signaling cascade (**Figure 3A**), suggesting that TLR signaling is activated in response oAβ42. It has been shown that inhibition of TLR signaling is sufficient to reduce the expression of microglial pro-inflammatory genes and secretion of pro-inflammatory cytokines in response to LPS and IL1β^73–75^. Importantly, direct interactions between amyloid peptides and TLR2 have been observed^76^. Microglia with increased expression of TLRs have been observed surrounding senile plaques in post-mortem human AD brains^77^, predominantly TLR2 and TLR4^78^. Thus, we asked whether TLR signaling was involved in mediating inflammatory responses specifically to oAβ42 in human microglia. We pre-treated iMGLs with either vehicle or a TLR inhibitor (TLRi) cocktail targeting TLR1, TLR2, and TLR4 for two hours before treating with oAβ42. Indeed, we saw a dramatic reduction in the expression pro-inflammatory genes, such as *IL1B*, *IL6*, and *TNF*, but not *NFKB1* in cells pre-treated with the TLRi (**Figure 3B-E**), demonstrating that TLR signaling is required to prompt a strong inflammatory response to oAβ42 in microglia. We also observed a downregulation of TREM2 upon oAβ42 exposure (**Figure 3F**) in control iMGLs, indicating that TREM2 may have a dynamic role in inflammation which can involve suppression in the setting of a strong inflammatory response, which has been described previously^79,80^. Additionally, we confirmed that, at baseline, expression of certain DAM genes was downregulated in KO iMGLs compared to controls, such as *GPNMB* and *CD9* (**Figure 3G-H**). Furthermore, pre-treatment with TLRi led to increased expression of these genes, in addition to *MITF*, a proposed master regulator of the DAM signature^44^ (**Figure 3I**). Similarly, we confirmed that genes corresponding to the homeostatic signature were downregulated in response to oAβ42, such as *P2RY12* and *CX3CR1*, confirming our previous results (**Additional File 5A**), although these were not significantly altered by TLRi pre-treatment (**Figure 3J-K**). These findings highlight that TLR signaling affects inflammatory gene expression and expression of TREM2 and TREM2-related genes. This is further supported by previous studies showing that TLR inhibition reduced inflammation, but also increased the expression of *Trem2* and *Arg1*^75^, shifting microglia towards a more anti-inflammatory phenotype. Additionally, NFκB regulates the production of microRNA-34a, which downregulates the expression of TREM2^81^, suggesting a possible regulatory mechanism between NFκB and TREM2. Thus, it is plausible that either downstream signaling components or common effectors of TLR signaling may have functions in regulating TREM2.

**Figure 3:**
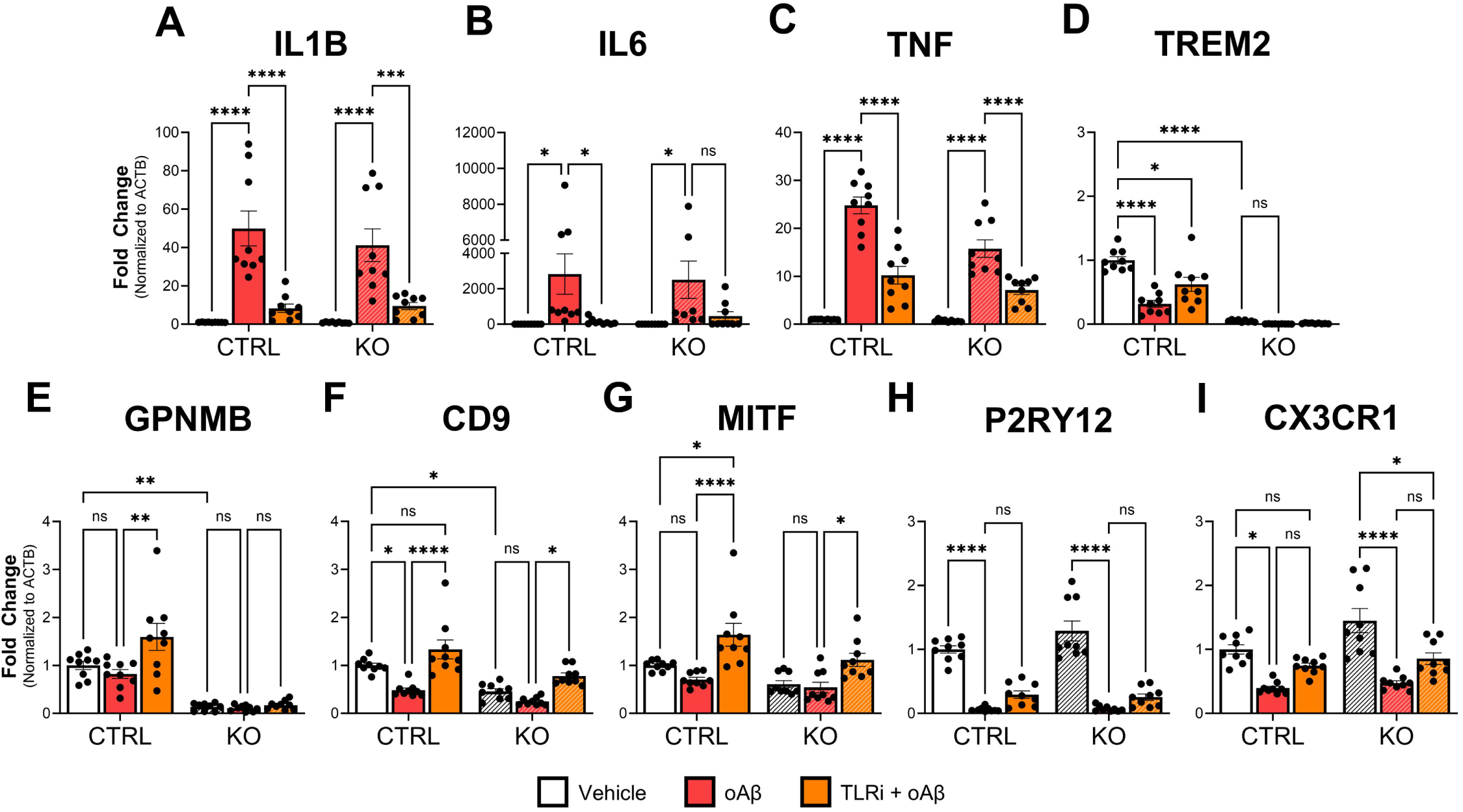
Inhibition of Toll-Like Receptor Signaling Drives Pro-Inflammatory Gene Expression in Response to Oligomeric Aβ42. **(A)** Schematic of TLR signaling involving NFκB activation, which leads to increased expression of pro-inflammatory genes. Created using BioRender. (**B-K**) qPCR analysis for a panel of pro-inflammatory (**B-E**), DAM (**F-I**), and homeostatic genes (**J-K**) in oAβ42-treated (red) and TLR inhibitor plus oAβ42-treated iMGLs (orange). Fold change was normalized to the control group (vehicle-treated CTRL iMGLs) and normalized to ACTB. Two-way ANOVA was used for statistical significance testing (* p-value ≤ 0.05, ** p-value ≤ 0.01, *** p-value ≤ 0.001, and **** p-value ≤ 0.0001), followed by Tukey’s post hoc analysis. Error bars are shown as SEM. Data was obtained from 3 independent experiments.

### 3.5 Secretion of cytokines in response to oligomeric amyloid-β are largely TREM2-independent

Next, we sought to determine if TREM2 expression affected cytokine production at the protein level in response to oAβ42. In line with our observations that pro-inflammatory processes are largely TREM2-independent, we found that the secretion of pro-inflammatory and anti-inflammatory cytokines and chemokines were similar between CTRL and KO iMGLs in response to (**Figure 4A-I**). oAβ42 treatment significantly drove the expression of 5 cytokines/chemokines assessed in our panel in both cell lines: IL6, TNFα, IL8, ILRA, and IL12-p40. IL1β production was increased in KO iMGLs in response to oAβ42 but was only trending in CTRL iMGLs. Surprisingly, the only cytokine that was significantly different between CTRL and KO iMGLs was the anti-inflammatory cytokine IL10, which was secreted at higher levels in KO iMGLs (**Figure 4E**). IL10 is traditionally thought of as an anti-inflammatory cytokine with critical roles in preventing inflammatory and autoimmune pathologies^82^, mainly acting as a feedback regulator that can act to regulate immune responses in innate immune cells^83^. However, increasing evidence suggests that IL10 may play a dual role in immunological homeostasis with one study showing that increased IL10 expression was associated with increase amyloid accumulation and impaired memory in APP mice^84^. Additionally, we observed a trend where secretion of IL1β was increased in KO iMGLs in response to oAβ42 relative to CTRL iMGLs (two-way ANOVA TREM2 effect p-value of 0.0512) (**Figure 4A**). Overall, these data show that TREM2 is not required for the secretion of inflammatory factors in response to oAβ42 in our model system, although it may reflect a more subtle role in balancing pro- and anti-inflammatory signals during the inflammatory cascade.

**Figure 4:**
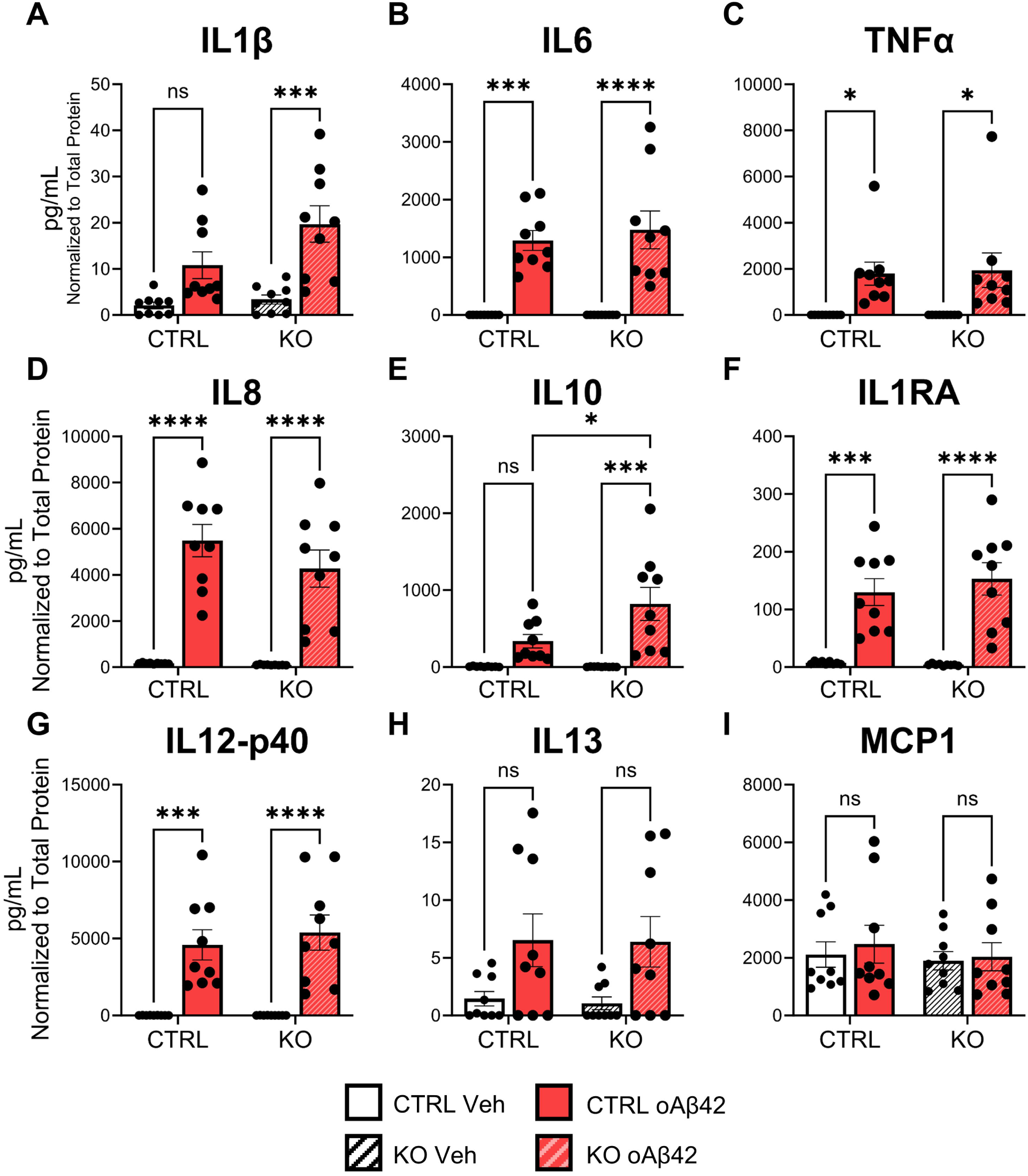
Quantification of Cytokines from iMGL Conditioned Media in Response to Oligomeric Aβ42. (**A-I**) Cytokine and chemokine quantification in iMGL conditioned media reported in pg/mL and normalized to the total protein in conditioned media. Secreted factors were quantified by Eve Technologies using a Luminex bead-based assay. Two-way ANOVA was used for statistical significance testing (* p-value ≤ 0.05, ** p-value ≤ 0.01, *** p-value ≤ 0.001, and **** p-value ≤ 0.0001), followed by Tukey’s post hoc analysis. Error bars are shown as SEM. Data was obtained from 3 independent experiments.

### 3.6 Altered lipid metabolism in TREM2 KO microglia

Given that TREM2 KO iMGLs showed elevated expression of cholesterol biosynthesis genes at baseline, and that lipid metabolism emerged as one of the most prominently downregulated pathways in response to oAβ42, we next asked whether there were lipid species with differential abundance between CTRL and KO iMGLs using targeted lipidomics from 3 independent differentiations (see raw and normalized lipidomics data in **Additional File 7**). We first looked broadly across major lipid classes and observed diverse changes at baseline and in response to oAβ42. Free cholesterol (FC) and cholesterol esters (CE) were increased in KO iMGLs at baseline compared to controls (**Figure 5A**), corroborating our transcriptomic findings. Under basal conditions, we observed a reduction of many lipid classes in KO iMGLs compared to controls, including bis(monoacylglycerol)phosphate (BMP), acylphosphatidylglycerol (AcylPG), lysophosphatidylcholine (LPC), lysophosphatidylethanolamine (LPE), lysophosphatidylinositol (LPI), phosphogylceride (PG), and phosphatidylserine (PS), in addition to specific sphingolipids, such as lactosylceramide (LacCer) and sphingomyelin (SM) (**Figure 5A**). We observed similar trends in BMP and cholesterol in our CRISPRi TREM2 KD model (**Additional File 8A**). When looking at the distribution of these lipids across samples, we see a modest increase in sterol lipids in KO and KD iMGLs (**Figure 5B; Additional File 8B**) and a subtle decrease in total sphingolipid molar percentages (**Figure 5B**). Overall, our data shows that loss of TREM2 in microglia leads to differences in lipid abundance, mainly in glycerophospholipids,sterol lipids, and sphingolipids, which may impact microglial membrane dynamics. Specifically, cholesterol esters and BMP levels are drastically different between controls and TREM2 KO iMGLs, which are associated with phagocytosis, lipid droplet biogenesis, and lysosomal function.

**Figure 5:**
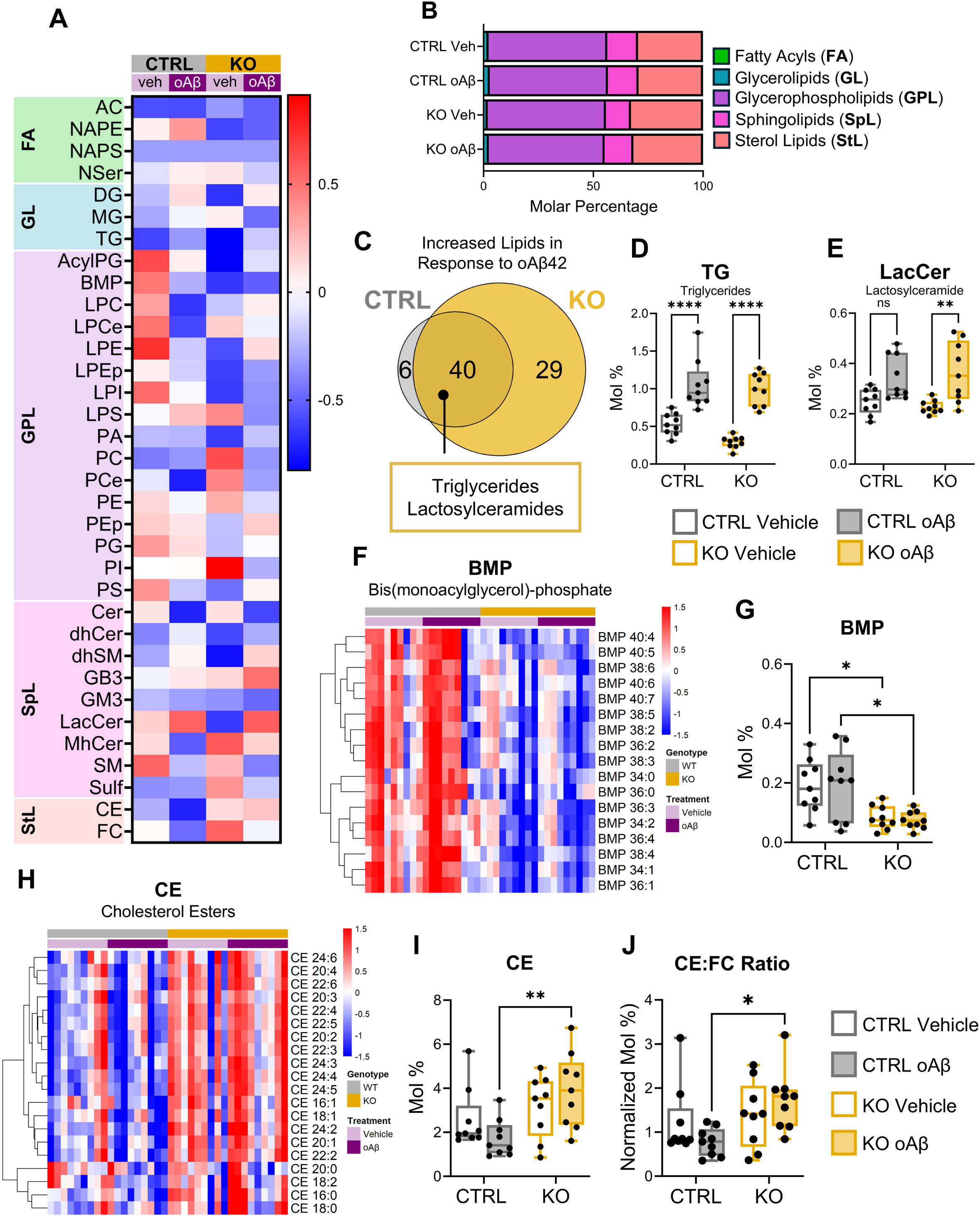
Lipidomic Changes in iMGLs at Steady-State and in Response to Oligomeric Aβ42. **(A)** Heatmap of lipid species showing the z-score of lipid concentration normalized to total protein. **(B)** Composition of lipids across conditions with the molar percentages of all lipid classes. (**C**) Venn diagram comparing significantly increased lipid species in CTRL and TREM2 KO iMGLs in response to oAβ42, with shared lipid classes highlighted in the box. (**D-E**) Boxplots showing the molar percentages for (**D**) TG and (**E**) and LacCer lipid species across all samples. (**F**) Heatmap of normalized z-score molar percentages for individual lipid species in the BMP class across all samples. (**G**) Boxplot showing the molar percentages for BMP. (**H**) Heatmap of normalized z-score molar percentages for individual lipid species in the CE class across all samples. (**I**) Boxplot showing the molar percentages for CE. (**J**) Boxplot showing the normalized molar percentages of the CE-to-free cholesterol (FC) ratio across all samples. Statistical analysis for boxplots was performed using two-way ANOVA was used for statistical significance testing, followed by Tukey’s post hoc analysis (* p-value < 0.05, ** p-value < 0.01, and **** p-value < 0.0001). Boxplots show data from the 25th to the 75th percentile, with the whiskers extending to the minimum and maximum values and the line in the box corresponding to the median. Data was obtained from 3 independent experiments.

We next asked how lipids changed in response to oAβ42 exposure. We performed differential abundance analysis across all lipid species in response to oAβ42 and found 46 lipid species that were significantly increased in controls and 69 lipid species that were significantly increased in KO iMGLs, with 40 of these lipidomic changes being shared between the two groups (**Figure 5C**). These shared lipid species that were increased consisted of primarily of triglycerides (TG) and LacCer (**Figure 5C**), which were also increased in response to oAβ42 in our KD model (**Additional File 8C**). When looking broadly at these lipid species, we observed a significant increase in the abundance of these lipids in response to oAβ42 in both CTRL and KO iMGLs (**Figure 5D-E**). LacCer is a bioactive lipid mediator that has been shown to promote inflammation via NFκB, suggesting that microglia may co-opt similar mechanisms to promote inflammation in response to oAβ42 via lipid alterations^85,86^.

We further assessed BMP and CE lipid species to better understand how these lipids were changing across conditions. We observed a significant reduction of multiple BMP species (**Figure 5F-G**) and increase in multiple CE species (**Figure 5H-I**) in KO iMGLs compared to controls, regardless of oAβ42 treatment, although CEs seem to accumulate more strongly in response to oAβ42 in KO iMGLs (**Figure 5I**). To assess changes in cholesterol trafficking, we looked at changes in the CE:FC ratio. We found that the CE:FC ratio was significantly increased in oAβ42-treated KO iMGLs, reflecting increased cholesterol trafficking in KO iMGLs (**Figure 5J**). While not significant, the CE:FC ratio nearly doubled in KO iMGLs at baseline compared to controls (**Figure 5J**), further suggesting that TREM2 KO iMGLs have increased cholesterol trafficking. These trends could also be observed in our TREM2 KD model (**Additional File 8F-I**). For BMP, we observed a significant reduction in BMP levels in TREM2 KO iMGLs (**Figure 5G**), suggesting that TREM2 expression affects the abundance of BMP in microglia. A similar significant reduction of BMP levels was observed in our KD model (**Additional File 8H**). Additionally, BMPs have also been shown to impact cholesterol trafficking, providing a link between BMP and cholesterol levels^87^. Although not significant, we found that CEs were increased almost two-fold in TREM2 KO iMGLs at baseline, which dramatically increased in KO iMGLs in response to oAβ42 (**Figure 5I**), a response not observed in isogenic controls. While we did not see a significant effect on CEs in our KD model, we did observe a significant increase in the abundance of free cholesterol (**Additional File 8J**).Similarly, we observed an increase in the CE:FC ratio in TREM2 KD iMGLs, although not significant (**Additional File 8K**). We next asked how TREM2 expression affects cholesterol trafficking. We grouped samples by genotype across treatments and cell lines and found that complete loss or reduction of TREM2 results in an increased CE:FC ratio, suggesting that TREM2 impacts cholesterol trafficking in microglia (**Additional File 8L**). Further studies are required to validate these potential changes in cholesterol trafficking experimentally. Recently, TREM2 was shown to regulate cholesterol metabolism under chronic phagocytic challenge using mouse models of MS^29^. In this demyelination mouse model, they found that loss of TREM2 led to decreased expression of cholesterol efflux genes, which may contribute to CE accumulation, which leads to impairments in phagocytosis^29^. Furthermore, increased CE levels have been linked with accumulation of lipid droplets (LDs) and subsequent impairments in phagocytosis^35,88^, emphasizing that microglia require tight regulation of cholesterol levels to efficiently enact their functions.

Taken together, we characterized changes in lipids in response to oAβ42 and identified BMP and CE as lipids that may be regulated in a TREM2-dependent manner. These observed changes likely have functional consequences in microglia that affect their responses to oAβ42 or impact their normal functions at baseline. Similarly to our transcriptomic data, we identified similar changes in lipid abundance between CTRL and KO iMGLs that are increased upon oAβ42 treatment, possibly reflecting lipid alterations that occur in response pathological oligomers. These data highlight the importance of TREM2 in maintaining lipid homeostasis.

## 4 DISCUSSION

DAM were first described in the 5xFAD mouse model, then subsequently described in other models, such as the APP-PS1^22^ and CK-p25 model^89^. This transcriptomic signature was not unique to Alzheimer’s disease, as it was also observed in mouse models of ALS and MS^21,22^. Importantly, over the course of disease, microglia were shown to transition from homeostatic microglia to DAM in a TREM2-dependent manner^21,22^. Comparison of mouse and human microglial transcriptomic studies have revealed that the DAM signature in mice does not correspond one-to-one with human microglial disease-associated signatures^27^, with murine DAM genes being expressed across many human microglial clusters^24–26^. One study in humans identified three distinct disease-associated clusters, which were associated with lipid metabolism, inflammation, and ribosome biogenesis^25^. Human microglia from the Sun study^24^ showed that DAM genes were primarily expressed in their lipid processing cluster MG4^24^. Here, we showed that there was a unique set of TREM2-dependent genes that were downregulated in TREM2 KO and KD iMGLs under steady-state conditions, which overlapped with DAM genes identified in mice^21^ (**Figure 1; Additional File 4**). Additionally, there was a significant downregulation of genes that corresponded to lipid-associated DAM and lipid-processing microglial signatures in TREM2 KO iMGLs. We observed a significant downregulation of these disease-associated genes at baseline in the absence of pathological exposure to oAβ42. While this iMGL model system is useful for studying human microglia in vitro, these cells are still relatively immature, which may explain why the disease associated signature is highly expressed at baseline, consistent with a previous study showing that the DAM signature was associated with embryonic microglia^90^. Importantly, we identified a unique set of TREM2-dependent genes that overlap with disease-associated signatures observed in mice and humans, providing further evidence that TREM2 is required for the maintenance of disease-associated genes in human microglia. Interestingly, we found that many of these disease-associated genes were associated with lipid metabolism, such as *GPNMB*, *MITF*, *PLA2G7*, and others, emphasizing a possible role of TREM2 in maintaining microglial lipid homeostasis.

Under steady-state conditions, we found a striking upregulation of genes related to cholesterol biosynthesis and it was the top pathway enriched in TREM2 KO iMGLs (**Figure 1B**). We also observed a similar trend in TREM2 KD iMGLs (**Additional File 4A**). We showed that cholesterol biosynthesis genes were upregulated in KO iMGLs, but cholesterol efflux genes were downregulated (**Figure 1C**). The expression of biosynthetic enzymes is regulated by the transcription factor sterol regulatory binding protein 2 (*SREBF2*), which was significantly upregulated in TREM2 KO iMGLs (**Figure 1C**), suggesting that dysregulation of cholesterol biosynthesis may be mediated by *SREBF2*, although further studies are required to better understand how TREM2 regulates SREBF2. Recently, it was shown that human microglia upregulate lipid metabolism genes in response to amyloid pathology, but is lost in human subjects with the TREM2 R62H variant^91^, strengthening the link between TREM2 and lipid metabolism. Excess cholesterol in cells is converted into CE and is stored within LDs or exported via efflux pathways. We found that cholesterol transporters, such as *ABCA1*, were downregulated in TREM2 KO iMGLs (**Figure 1C**). These changes in cholesterol trafficking highlight the importance of maintaining cholesterol homeostasis in microglia by balancing synthesis and export. Cholesterol contributes to membrane rigidity in cells and is concentrated in lipid rafts, which are specialized regions in the plasma membrane that promotes the recruitment of membrane proteins and acts as a scaffold for signaling molecules^92^. Furthermore, over 250 cholesterol-binding proteins have been identified, with many being genetically associated with neurologic disease^93^. Thus, alterations in cholesterol metabolism may affect diverse processes in microglia which affect their function in disease. This points towards TREM2 as a possible regulator of cholesterol metabolism in microglia. Previous studies have implicated TREM2 in cholesterol metabolism, showing that loss of *Trem2* led to increased CE levels in a mouse model of demylination^29^. Similar increases in CE were observed in APOE KO iMGLs, suggesting an interplay between TREM2 and APOE^29^. Here, we showed that loss of TREM2 also led to a downregulation of *APOE* at baseline (**Figures 1C**), which may further exacerbate lipid accumulation in microglia. Interestingly, in response to oAβ42, we found that accumulation of CE was higher in TREM2 KO iMGLs (**Figure 5I**), mirroring similar responses observed in demyelination models.

The esterification of cholesterol is mediated by acetyl-CoA/sterol-O acetyltransferases (ACAT/SOAT) and lecithin-cholesterol acyltransferase (LCAT).Inhibition of ACAT in iMGLs was shown to improve phagocytosis by modulating expression of LRP1^35^, providing an important mechanistic link between cholesterol homeostasis and regulation of phagocytosis. While many studies have shown that mouse and human TREM2 KO microglia exhibit impaired phagocytosis^62,68–70^, increasing evidence suggests that these impairments may be a result of dysfunctional cholesterol trafficking^35,94^. Membrane rigidity and fluidity are carefully balanced and regulated through diverse lipids, including glycerophospholipids, sphingolipids, and sterols^37^. Given that phagocytosis requires extensive membrane remodeling and cholesterol inhibits membrane curvature, it is possible that excess cholesterol could impair phagocytosis. This provides a possible mechanistic link between cholesterol and phagocytic impairments, although further work is required to better understand this process. These studies highlight that cholesterol imbalance leads to dysregulated microglial function.

In addition to cholesterol dysregulation, we observed a consistent reduction of BMP in TREM2 KO iMGLs under both vehicle and oAβ42-treated conditions (**Figure 5F, H**). BMPs are located on the surface of intraluminal vesicles within lysosomes and function primarily to stimulate lysosomal lipid catabolism. It has been extensively studied in the context of Niemann-Pick type C disease, a lysosomal storage disorder, where BMP accumulates in the lysosome and impairs cholesterol export from the lysosome leading to neurodegeneration^95,96^. Niemann-Pick C1 and C2 (NPC1 and NPC2) are lysosomal cholesterol transporters that rely on BMP for their normal function and loss of function mutations in NPC1 and NPC2 are causative for Niemann-Pick type C disease^97^. One study found that reduction of BMP led to gangliosidosis and lysosomal abnormalities^98^. In the absence of TREM2, we observed a strong depletion of BMP across all conditions (**Figure 5F, H**), which may lead to lysosomal dysfunction and inefficient phagocytic processing. Taken together, this highlights the significant need for functional studies that show how altered BMP levels affect lysosomal function and phagocytosis.

In response to oAβ42, we observed an increase in TG and LacCer (**Figure 5C-E**), suggesting that TGs may be involved in the immune response to oAβ42. Diacylglycerol O-acyltransferases (DGATs) and long-chain acyl-CoA synthetases (ACSLs) are critical enzymes involved in the generation of acyl-CoA, which is required for esterification of cholesterol and production of TG. ACSL1 has been shown to be a critical enzyme involved in the formation of LDs, with increased expression in LD-rich microglia and AD patients^88^. Importantly, LD formation was influenced by APOE genotype, where APOE4 affects lipid metabolic shifts in microglia^99^. These data further show that APOE also contributes significantly to lipid dysregulation in microglia, in addition to TREM2. Additionally, TG synthesis requires glycerol and fatty acid chains, which are both generated during glycolysis. Transcriptomic analysis revealed that glycolytic genes were strongly upregulated in response to oAβ42 in both CTRLs and KO iMGLs (**Additional File 5H**). Therefore, increased glycolysis could potentially contribute to increased TG production in response to oAβ42. Our findings provide a possible link between increased LDs and glycolysis as a common feature during strong pro-inflammatory responses. In support of this, treatment of iMGLs with LPS resulted in accumulation of TG-rich LDs, where TG synthesis is essential for activation-induced transcription and secretion of inflammatory cytokines and chemokines^100^. TGs appear to be increased in iMGLs in response to oAβ42, likely promoting LD formation and supporting inflammation while impairing phagocytosis.

We also observed increased levels of LacCer in response to oAβ42 (**Figure 5C, E**). LacCer is a bioactive lipid that responds to specific stimuli and is involved in signaling pathways. Additionally, LacCer is found on lipid rafts and supports inflammatory signaling^101^. In regards to inflammation, LacCer has been shown to interact and activate cytosolic phospholipase A2 (cPLA2), which leads to downstream activation of NFκB^102^ and can regulate gene expression of inducible nitric oxide synthase through Ras/ERK and NFκB pathways^85^, supporting a role for LacCer in supporting inflammatory processes. Thus, increased LacCer following oAβ42 exposure may represent another lipid-mediated mechanism through which inflammatory signaling is amplified independently of TREM2.

Interestingly, we observed a downregulation of TREM2 in response to oAβ42 (**Figure 3F**), which was accompanied by the upregulation of genes associated with pro-inflammatory signaling pathways. Similarly, a study showed that treatment of primary murine microglia with LPS led to downregulation of *Trem2*^79^, suggesting that lower TREM2 may be required to mount a strong pro-inflammatory response. Our analysis revealed that a subset of inflammatory genes related to chemokine and cytokine signaling were more highly expressed in TREM2 KO iMGLs (**Figure 2G**), which may further indicate that loss of TREM2 promotes inflammation. Inhibition of TLR signaling led to a significant downregulation of pro-inflammatory genes (**Figure 3B-D**), suggesting that TLR signaling, not TREM2, is driving the observed inflammatory response in microglia. While *TREM2* was downregulated in response to oAβ42, pre-treatment with a TLR inhibitor cocktail led to an increase in *TREM2* gene expression in isogenic control iMGLs. Furthermore, TLR signaling inhibition led to significantly increased expression of TREM2-dependent genes, such as *GPNMB*, *CD9*, and *MITF*, in response to oAβ42 (**Figure 3G-I**). It has previously been shown that expression of NFκB can promote the production of microRNAs that lead to downregulation of TREM2^81^. Additionally, overexpression of TREM2 reduced inflammation by downregulating NFκB signaling^80^. These data suggest that TREM2 and NFκB signaling may negatively regulate each other to modulate inflammatory responses.

We identified CTRL- and TREM2 KO-specific pathways that were uniquely upregulated upon oAβ42 stimulation (**Additional File 6**). In CTRL iMGLs only, we observed upregulation of pathways associated with mitochondrial RNA (mtRNA) degradation (**Additional File 6A-B**). Mitochondrial DNA (mtDNA) and mtRNA are released under high stress conditions, which drives SASP^103^. Additionally, mtDNA activates cGAS signaling to promote inflammation^104^. These studies show that mtDNA and mtRNA can act as damage associated molecular patterns to promote inflammation. The activation of mtRNA degradation in CTRL iMGLs in response to oAβ42 suggests that inflammatory responses in microglia trigger mitochondrial stress. In TREM2 KO iMGLs, we observed an upregulation of genes associated with adipogenesis and SASP (**Additional File 6C-D**). Adiposomes in microglia promote inflammation, suggesting that loss of TREM2 may enhance inflammatory signaling via metabolic changes in lipids. Indeed, it has been reported that genetic perturbation of sphingolipid metabolism led to alterations in membrane lipids, which in turn affected TLR-driven production of IL6^37^, further strengthening the link between inflammatory signaling and lipid metabolism. Additionally, epigenetic regulation of adipogenesis was a significant pathway found in TREM2 KO iMGLs only (**Additional File 6C**), suggesting that epigenetic reprogramming may contribute to the changes in lipid metabolism observed. We observed increased expression of the transcription factor CREBBP only in TREM2 KO iMGLs (**Additional File 6D**), which has been shown to regulate lipid metabolism in microglia^105^. We also observed upregulation of genes associated with SASP, which is a common phenotype observed in senescent cells consisting of metabolic alterations and increased secretion of various cytokines^106^. This is accompanied by the recruitment of immune cells to eliminate senescent cells, but also promotes inflammation by effector cells. Importantly, AP1, which consists of cFOS and cJUN, can promote cellular senescence, which we show is upregulated uniquely in TREM2 KO iMGLs (**Additional File 6D**). This suggests that loss of TREM2 may drive low-level inflammation that accelerates senescence and impairs normal microglial functions upon stimulation with oAβ42.

A significant limitation of this current study is that the synthetic oAβ42 used for treatments does not reflect pathological Aβ plaques that are observed in AD, thus microglial responses *in vivo* may differ. However, oligomeric Aβ42 species are some of the earliest species generated during AD pathogenesis and oligomers have been found to surround senile plaques in AD^107^. Thus, using oligomers still proves useful in understanding some of the earliest responses to pathological changes that occur in AD. Xenotransplantation of human iPSC microglia represents a suitable alternative, where *in vivo* responses to AD pathology can be studied. These approaches are especially valuable when xenografting is performed in humanized AD mouse models, enabling the characterization of human microglial responses to human Aβ pathology. Another limitation of this study is that the responses described here are acute, short-term microglial responses, which do not capture the microglial changes that arise from chronic or long-term exposure to Aβ. Future studies incorporating microglial temporal dynamics would be useful in understanding how microglial responses to Aβ change over time.

## 5 CONCLUSIONS

Collectively, our findings support a model where TREM2 is not required for the initiation of acute inflammatory responses to oAβ42 but instead supports the metabolic and transcriptional programs necessary to maintain normal microglial homeostasis. Loss of TREM2 disrupted lipid-processing, altered cholesterol homeostasis, and impaired expression of lipid-associated DAM genes, suggesting that TREM2 primarily functions to coordinate metabolic changes, rather than directly driving inflammatory activation in our model. TREM2 is a complex regulator of diverse microglial processes and we highlight here how lipids may be involved in multiple facets of microglial responses, underscoring their relevance in AD.

## DATA AVAILABILITY

## FUNDING

C.L.C. was supported by a Diversity Fellowship from the NIH National Institute on Aging associated with the parent grant from A.F.T. (RO1AG073360). A.A.S. is supported by the Henry and Marilyn Taub Foundation and Thompson Family Foundation (TAME-AD). The Columbia Biomarkers Core Laboratory was supported by the NIH National Center for Advancing Translational Sciences (UL1TR001873 and UM1TR006153). The content is solely the responsibility of the authors and does not necessarily represent the official views of the NIH.

## AUTHOR CONTRIBUTIONS

C.L.C. performed all experiments, analyzed data, and wrote the manuscript. R.P. assisted with experiments and provided experimental feedback. G.M.F. and T-W.K. provided the FA10 TREM2 KO and isogenic control hiPSCs. A.A.S. and A.F.T. conceptualized and funded the study. All authors reviewed the manuscript.

## ACKNOWLEDGEMENTS

We would like to thank the Michael Kissner and Juan Idiarte from the Columbia University Stem Cell Initiative Flow Cytometry Core for their assistance with HPC validation and iMGL characterization experiments. We would also like to thank Dr. Alejandro Chavez and his lab for kindly providing us with the CRISPRi construct and for his assistance with generating TREM2 gRNA sequences for our CRISPRi TREM2 KD model. Lastly, we are thankful to Dr. Renu Nandakumar and Alexandra Dombroski from the Columbia University Biomarkers Core Laboratory for their assistance and guidance on performing and processing samples for targeted lipidomics, and to Dr. Estela Area-Gomez for helpful discussions regarding the interpretation of those results.

## LIST OF ADDITIONAL FILES

**Additional File 1.pptx: Quality Control of iMGLs and Validation of Experimental Model.**

(**A-F**) qPCR analysis showing expression of hiPSC markers (**A-B**) and microglial markers (**C-F**) in hiPSCs and differentiated microglia (iMGLs). Fold change was calculated using the 2^-ΔΔCT^ method and normalized against ACTB and hiPSC as the control. F-test for normality tests were performed to assess variance and normality, respectively. Student’s t-test was used for statistical significance testing (*** p-value ≤ 0.001 and **** p-value ≤ 0.0001).Error bars are shown as SEM. (**G**) Flow cytometry analysis of iMGLs positive for CD45 (y-axis) and CD11b (x-axis), canonical microglial markers used to isolate microglia *in vivo*^207^. (**H-I**) Immunofluorescence staining against IBA (**H**) and P2RY12 (**I**) in iMGLs. (**J**) Non-reducing SDS-PAGE of oAβ42 run in technical triplicate and stained with 6E10 amyloid antibody to validate the formation of oligomers used for experiments. (**K**) Western blotting of TREM2 in CTRL and TREM2 KO iMGLs to validate reduced TREM2 protein expression in iMGLs with GAPDH as housekeeping control. Data shown from one representative experiment in technical triplicate.

**Additional File 2.pptx: RNA-Sequencing Quality Control.**

(**A**) MA plot showing individual sample variance all others in the same experimental group. This plot highlights one sample (CTRL5), which had above average deviation compared to other vehicle controls. (**B**) The median absolute deviation differences were plotted for every sample, showing that CTRL5 (red) had above average deviation from the median compared to all other samples. (**C**) Principal component analysis of normalized and batch-corrected sequencing data across genotypes, treatment, and batch. Sequencing was performed in three independent batches for three biological replicates.

**Additional File 3.xlsx: DEGs and Pathway Analysis Results from Transcriptomics.**

Excel file containing the raw DEGs from both FA10 TREM2 KO and IMR90 TREM2 KD RNA-seq across multiple groups (steady-state, CTRL-response, KO-response). Pathway analysis results from the study. Data relates to Figure 1, Figure 2, Additional File 4, and Additional File 6.

**Additional File 4.pptx: Analysis of CRISPRi TREM2 KD iMGLs at Steady-State and in Response to Oligomeric Aβ42.**

(**A**) Pathway analysis results using GSEA to highlight reactome pathways that were enriched in iMGLs treated with NT gRNA (beige) or TREM2-targeting gRNA (orange) at steady-state. Pathways shown were selected from the top 50 significant pathways ranked by normalized enrichment score (NES). Pathways shown had a FDR less than 0.05. (**B**) Heatmap visualization of cholesterol biosynthesis and cholesterol efflux genes. (**C**) Heatmap visualization of DAM genes identified in mice, xenotransplanted iMGLs, and human microglial lipid-associated DAM genes (Lipo DAM). For heatmaps, normalized z-score gene expression values were used for visualization. (**D-E**) Venn diagrams comparing gene expression changes in response to oAβ42 in cells treated with non-targeting sgRNA (NT) or TREM2-targeting sgRNA (TREM2 KD). Significant upregulated (**D**) and downregulated (**E**) genes were used for visualization, those with a FDR < 0.05. Data was obtained from 1 experiment.

**Additional File 5.pptx: Gene Set Score Analysis of Human AD Microglial Signatures in iMGLs.**

(**A-L**) Boxplots showing gene set score analysis comparing human microglial gene signatures observed in AD and controls (Sun *et al*). (**M-O**) Boxplots showing gene set score analysis comparing microglial DAM subpopulations from the human microglial atlas (Martins-Ferreira *et al*). (**P**) Boxplot showing gene set score analysis of the DAM signature identified from xenotransplanted iPSC-derived microglia (xMGL) into the humanized *APP^NL-G-F^* mouse model. Two-way ANOVA was used for statistical significance testing followed by Tukey’s post hoc analysis (* p-value ≤ 0.05, ** p-value ≤ 0.01, *** p-value ≤ 0.001, and **** p-value ≤ 0.0001). Boxplots show data from the 25th to the 75th percentile, with the whiskers extending to the minimum and maximum values and the line in the box corresponding to the median.

**Additional File 6.pptx: CTRL and TREM2 KO-Specific Responses to Oligomeric Aβ42.**

(**A**) Pathway analysis of genes upregulated only in CTRL iMGLs in response to oAβ42 using Reactome pathways, with the dashed line showing the threshold for significance (FDR ≤ 0.05) and the color corresponding to the gene ratio. (**B**) Heatmap of normalized z-score expression of genes relating to mitochondrial RNA degradation, which were extracted from the corresponding pathways. (**C**) Pathway analysis of genes upregulated only in TREM2 KO iMGLs in response to oAβ42 using Reactome pathways. (**D**) Heatmap of normalized z-score expression of genes that are part of the corresponding pathways, annotated on the left of the heatmap.

**Additional File 7.xlsx:Lipidomics Molar Percentages and Normalized Concentration.**

Excel file containing the raw molar percentages and normalized lipid concentrations for targeted lipidomics performed in FA10 TREM2 KO and IMR90 TREM2 KD iMGLs.

**Additional File 8.pptx: Lipidomics of CRISPRi TREM2 KD iMGLs at Steady-State and in Response to Oligomeric Aβ42.**

(A) Heatmap of lipid species showing the z-score of lipid concentration normalized to total protein. (B) Composition of lipids across conditions with the molar percentages of all lipid classes. (**C**) Venn diagram comparing significantly increased lipid species in non-targeting control iMGLs and TREM2 KD iMGLs in response to oAβ4242, with shared lipid classes highlighted in the box. (**D-E**) Boxplots showing the molar percentages for increased lipid species across all samples. Two-way ANOVA was used for statistical significance testing, followed by Tukey’s post hoc analysis (* p-value < 0.05 and ** p-value ≤ 0.01). Error bars are shown as SEM. (**F-G**) Heatmap of normalized z-score molar percentages for individual lipid species for BMP (**F**) and CE (**G**) across all samples. (**H-J**) Boxplots showing the molar percentages for BMP (**H**), cholesterol esters (**I**), and free cholesterol (**J**) across all samples. Two-way ANOVA was used for statistical significance testing (* p-value < 0.05) with p-value reported for the genotype effect (gRNA). Error bars are shown as SEM. (**K**) Boxplot showing the normalized molar percentages of the CE-to-free cholesterol (FC) ratio across all samples. (**L**) Boxplot showing the normalized molar percentages of the CE-to-FC ratio across all samples from both hiPSC lines grouped by genotype (CTRL = FA10 CTRL and IMR90 NT, KO/KD = FA10 TREM2 KO and IMR90 TREM2 KD). Statistical analysis for boxplots was performed using Mann Whitney test for non-parametric data (*** p-value < 0.001). Data was obtained from 1 experiment.

**Additional File 9.xlsx: List of Primer Sequences for TREM2 gRNA Cloning and qRT-PCR.**

Two sheets providing oligonucleotide sequences that were designed and used for TREM2 gRNA cloning and qRT-PCR.

## LIST OF ABBREVIATIONS

KO: knockout
oAβ42: oligomeric amyloid-β 42
Aβ: amyloid-β
APP: amyloid precursor protein
AD: Alzheimer’s disease
scRNA-seq: single-cell RNA sequencing
snRNA-seq: single-nucleus RNA sequencing
DAM: disease-associated microglia
iPSC: induced pluripotent stem cells
TLR: toll-like receptor
iMGL: iPSC-derived microglia-like cells
CTRL: isogenic control
RNA-seq: bulk RNA sequencing
DEG: differentially expressed gene
CRISPRi: clustered regularly interspaced short palindromic repeats inhibition
gRNA: guide
RNA NT: non-targeting
KD: knockdown
HuMicA: human microglial atlas
xMGL: xenotransplanted iPSC-derived microglia
Lipo DAM: lipid-associated DAM
PCA: principal component analysis
BMP: bis(monoacylglycero)-phosphate
LPC: lysophosphatidylcholine
LPE: lysophosphatidylethanolamine
LacCer: lactosylceramide
FC: free cholesterol
CE: cholesterol ester
TG: triglyceride
LD: lipid droplet
LXR: liver X receptor
RXR: retinoid X receptor
ACAT: acyl-CoA acetyltransferase
NPC: Niemann-Pick disease type C
ACSL: acyl-CoA synthetase long-chain
mtDNA: mitochondrial DNA
mtRNA: mitochondrial RNA
SASP: senescence-associated secretory phenotype

## Notes

### Competing Interest Statement

The authors have declared no competing interest.

