## Additional File 1 for "Transcriptomic and Lipidomic Profiling of Human iPSC-Derived Microglia Reveals TREM2-Dependent and -Independent Processes"

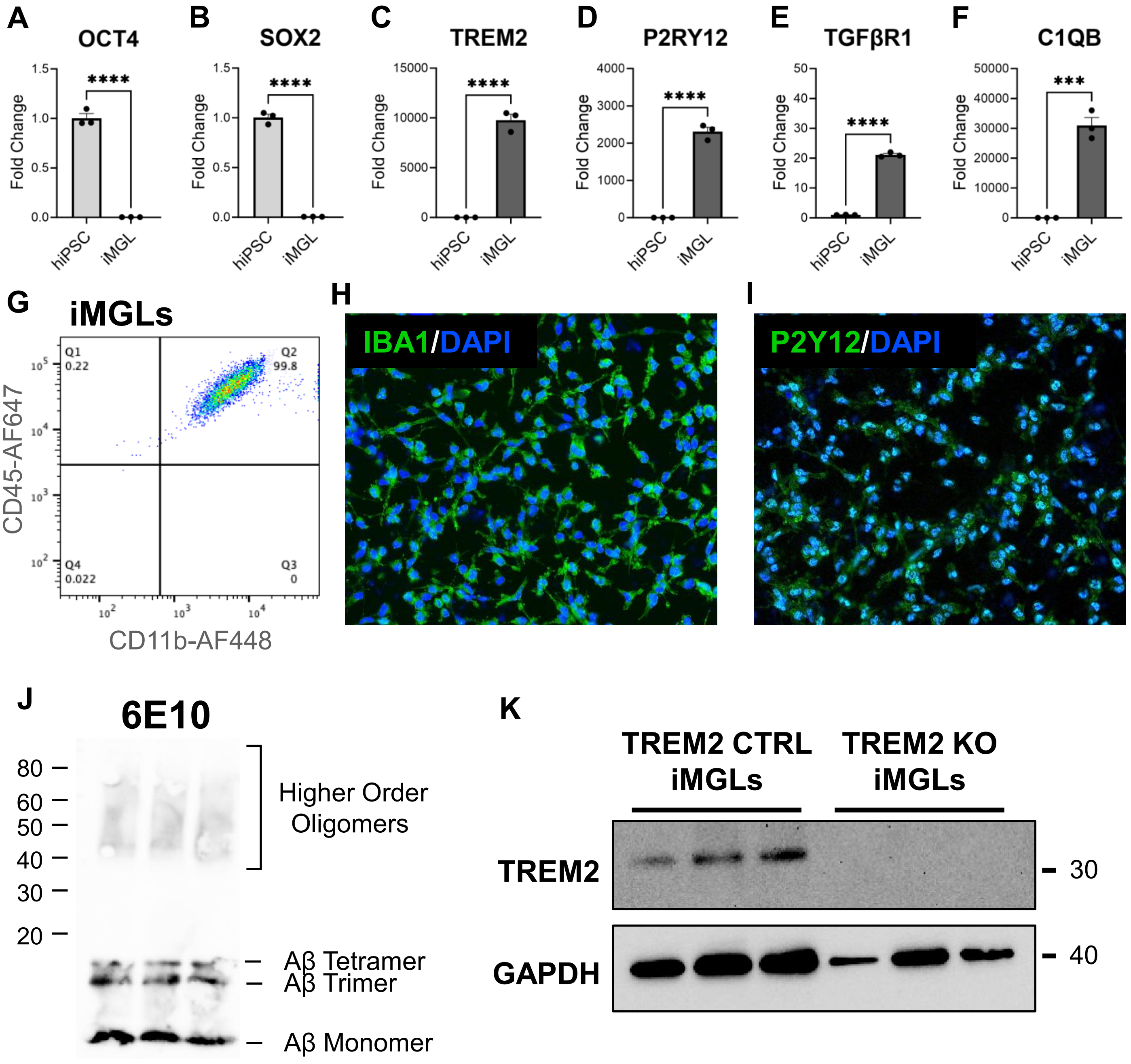


**Additional File 1: Quality Control of iMGLs and Validation of Experimental Model.**
(**A-F**) qPCR analysis showing expression of hiPSC markers (**A-B**) and microglial markers (**C-F**) in hiPSCs and differentiated microglia (iMGLs). Fold change was calculated using the $2^{-\Delta\Delta CT}$ method and normalized against ACTB and hiPSC as the control. F-test for normality tests were performed to assess variance and normality, respectively. Student's t-test was used for statistical significance testing (*** p-value ≤ 0.001 and **** p-value ≤ 0.0001).Error bars are shown as SEM. (**G**) Flow cytometry analysis of iMGLs positive for CD45 (y-axis) and CD11b (x-axis), canonical microglial markers used to isolate microglia *in vivo*[207]. (**H-I**) Immunofluorescence staining against IBA (**H**) and P2RY12 (**I**) in iMGLs. (**J**) Non-reducing SDS-PAGE of oAβ42 run in technical triplicate and stained with 6E10 amyloid antibody to validate the formation of oligomers used for experiments. (**K**) Western blotting of TREM2 in CTRL and TREM2 KO iMGLs to validate reduced TREM2 protein expression in iMGLs with GAPDH as housekeeping control. Data shown from one representative experiment in technical triplicate.
