## Additional File 2 for "Transcriptomic and Lipidomic Profiling of Human iPSC-Derived Microglia Reveals TREM2-Dependent and -Independent Processes"

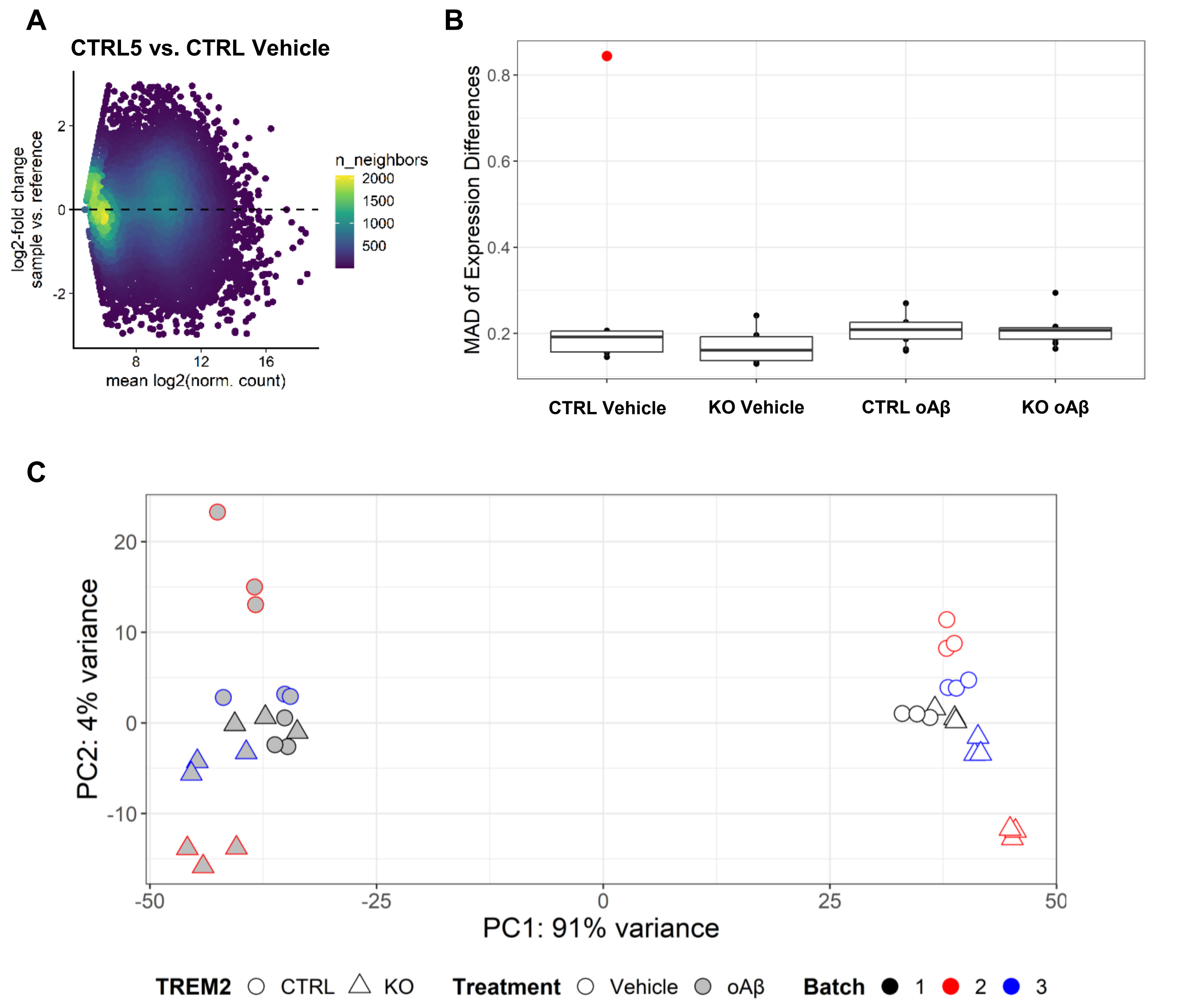


**Additional File 2: RNA-Sequencing Quality Control.**
(**A**) MA plot showing individual sample variance all others in the same experimental group. This plot highlights one sample (CTRL5), which had above average deviation compared to other vehicle controls. (**B**) The median absolute deviation differences were plotted for every sample, showing that CTRL5 (red) had above average deviation from the median compared to all other samples. (**C**) Principal component analysis of normalized and batch-corrected sequencing data across genotypes, treatment, and batch. Sequencing was performed in three independent batches for three biological replicates.
