## Additional File 4 for "Transcriptomic and Lipidomic Profiling of Human iPSC-Derived Microglia Reveals TREM2-Dependent and -Independent Processes"

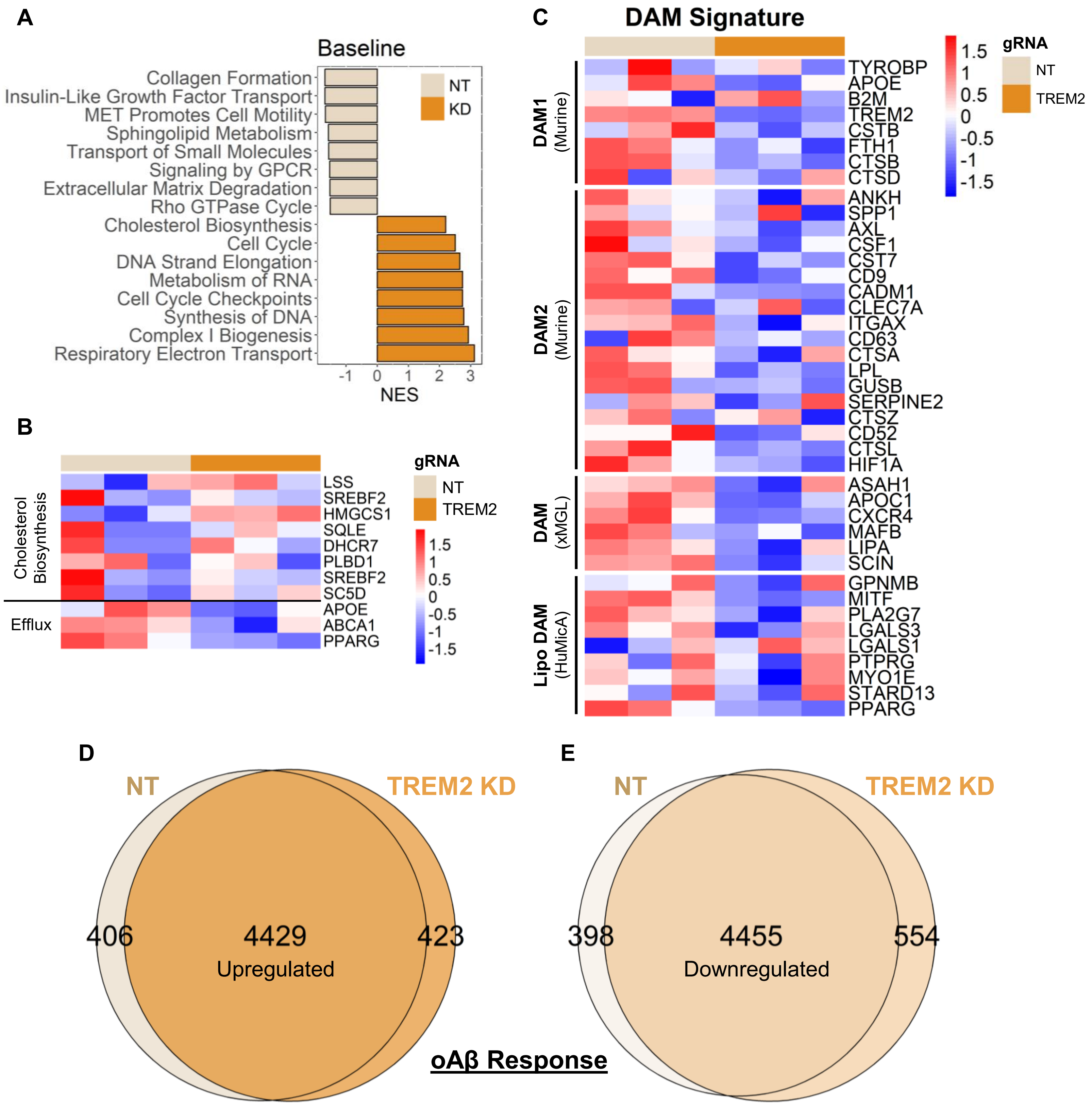


**Additional File 4: Analysis of CRISPRi TREM2 KD iMGLs at Steady-State and in Response to Oligomeric Aβ42.**

(**A**) Pathway analysis results using GSEA to highlight reactome pathways that were enriched in iMGLs treated with NT gRNA (beige) or TREM2-targeting gRNA (orange) at steady-state. Pathways shown were selected from the top 50 significant pathways ranked by normalized enrichment score (NES). Pathways shown had a FDR less than 0.05. (**B**) Heatmap visualization of cholesterol biosynthesis and cholesterol efflux genes. (**C**) Heatmap visualization of DAM genes identified in mice, xenotransplanted iMGLs, and human microglial lipid-associated DAM genes (Lipo DAM). For heatmaps, normalized z-score gene expression values were used for visualization. (**D-E**) Venn diagrams comparing gene expression changes in response to oAβ42 in cells treated with non-targeting sgRNA (NT) or TREM2-targeting sgRNA (TREM2 KD). Significant upregulated (**D**) and downregulated (**E**) genes were used for visualization, those with a FDR < 0.05. Data was obtained from 1 experiment.
