## Additional File 5 for "Transcriptomic and Lipidomic Profiling of Human iPSC-Derived Microglia Reveals TREM2-Dependent and -Independent Processes"

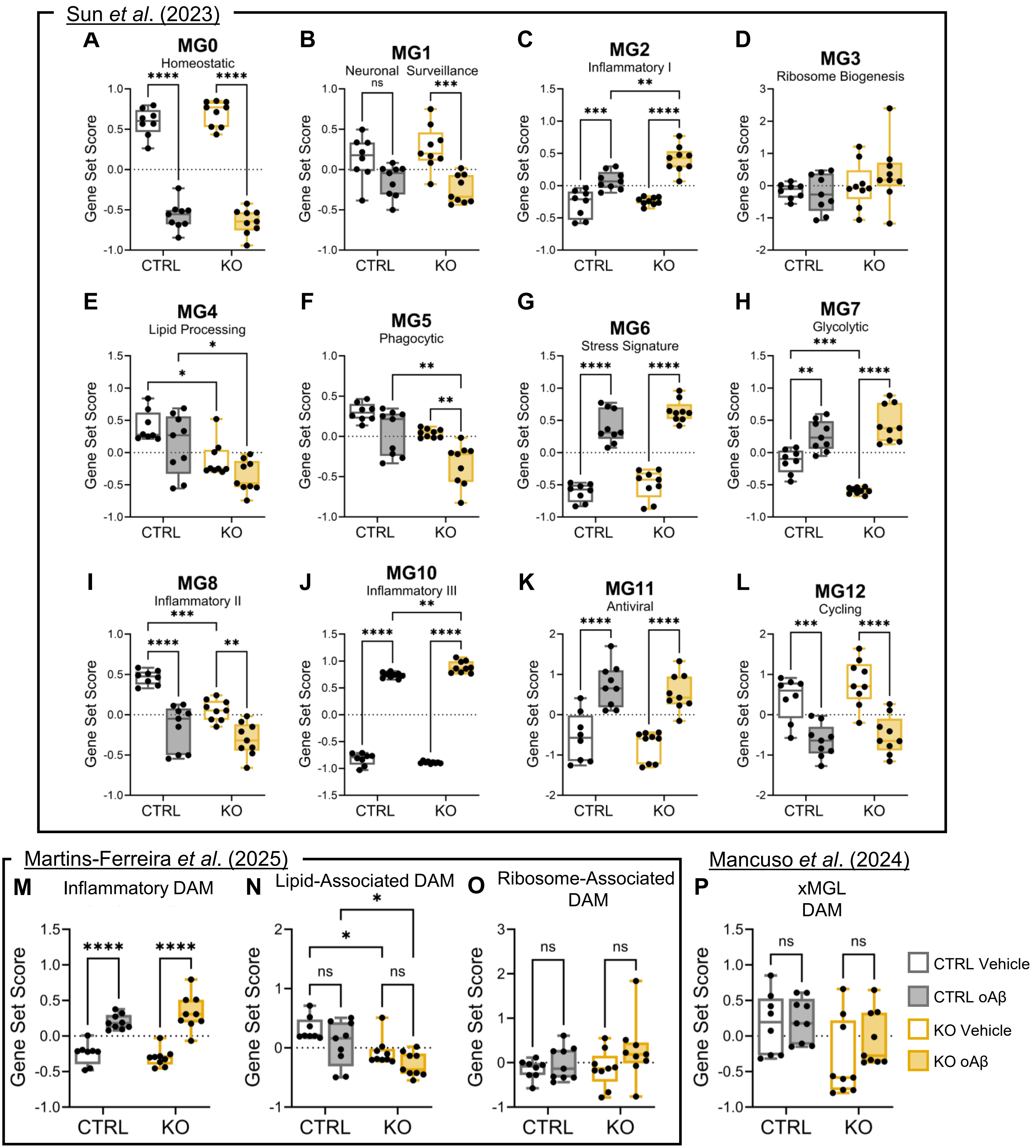


**Additional File 5: Gene Set Score Analysis of Human AD Microglial Signatures in iMGLs**.
(**A-L**) Boxplots showing gene set score analysis comparing human microglial gene signatures observed in AD and controls (Sun *et al*). (**M-O**) Boxplots showing gene set score analysis comparing microglial DAM subpopulations from the human microglial atlas (Martins-Ferreira *et al*). (**P**) Boxplot showing gene set score analysis of the DAM signature identified from xenotransplanted iPSC-derived microglia (xMGL) into the humanized *APP*$^{NL\text{-}G\text{-}F}$ mouse model. Two-way ANOVA was used for statistical significance testing followed by Tukey's post hoc analysis (* p-value ≤ 0.05, ** p-value ≤ 0.01, *** p-value ≤ 0.001, and **** p-value ≤ 0.0001). Boxplots show data from the 25th to the 75th percentile, with the whiskers extending to the minimum and maximum values and the line in the box corresponding to the median.
