## Additional File 6 for "Transcriptomic and Lipidomic Profiling of Human iPSC-Derived Microglia Reveals TREM2-Dependent and -Independent Processes"

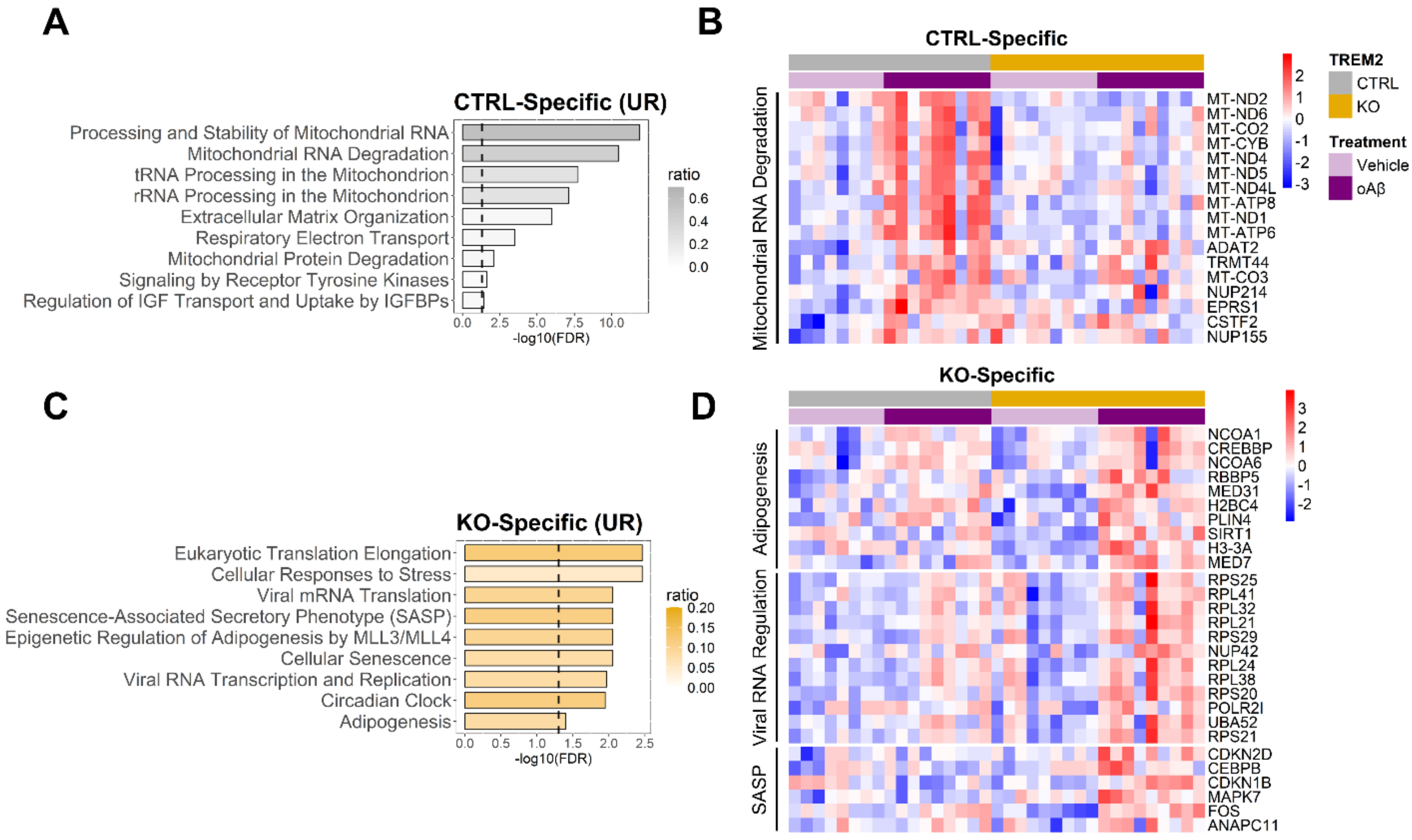


**Additional File 6: CTRL and TREM2 KO-Specific Responses to Oligomeric Aβ42**.
(**A**) Pathway analysis of genes upregulated only in CTRL iMGLs in response to oAβ42 using Reactome pathways, with the dashed line showing the threshold for significance (FDR ≤ 0.05) and the color corresponding to the gene ratio. (**B**) Heatmap of normalized z-score expression of genes relating to mitochondrial RNA degradation, which were extracted from the corresponding pathways. (**C**) Pathway analysis of genes upregulated only in TREM2 KO iMGLs in response to oAβ42 using Reactome pathways. (**D**) Heatmap of normalized z-score expression of genes that are part of the corresponding pathways, annotated on the left of the heatmap.
