## Additional File 8 for "Transcriptomic and Lipidomic Profiling of Human iPSC-Derived Microglia Reveals TREM2-Dependent and -Independent Processes"

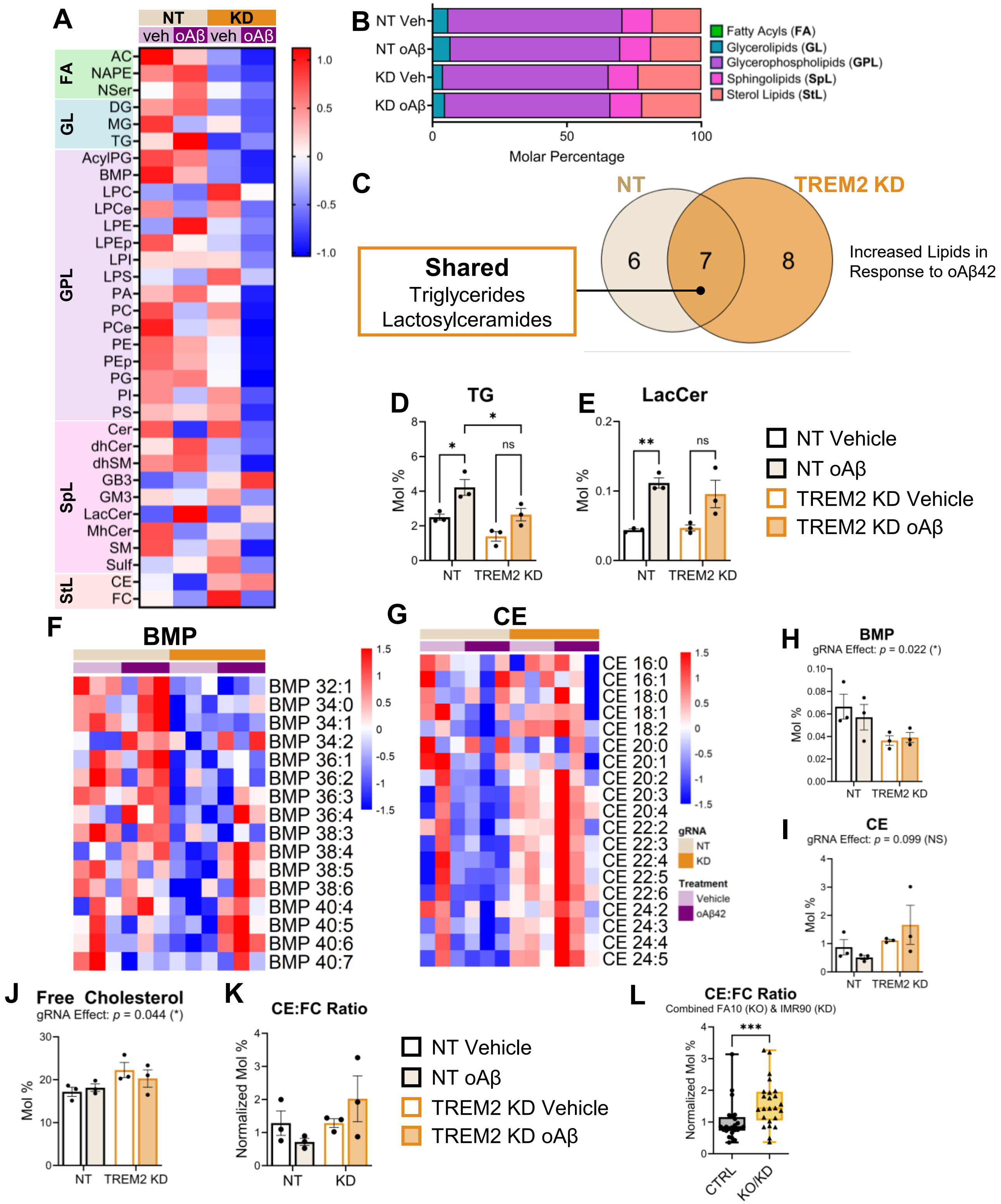


**Additional File 8: Lipidomics of CRISPRi TREM2 KD iMGLs at Steady-State and in Response to Oligomeric Aβ42**. (**A**) Heatmap of lipid species showing the z-score of lipid concentration normalized to total protein. (**B**) Composition of lipids across conditions with the molar percentages of all lipid classes. (**C**) Venn diagram comparing significantly increased lipid species in non-targeting control iMGLs and TREM2 KD iMGLs in response to oAβ4242, with shared lipid classes highlighted in the box. (**D-E**) Boxplots showing the molar percentages for increased lipid species across all samples. Two-way ANOVA was used for statistical significance testing, followed by Tukey's post hoc analysis (* p-value < 0.05 and ** p-value ≤ 0.01). Error bars are shown as SEM. (**F-G**) Heatmap of normalized z-score molar percentages for individual lipid species for BMP (**F**) and CE (**G**) across all samples. (**H-J**) Boxplots showing the molar percentages for BMP (**H**), cholesterol esters (**I**), and free cholesterol (**J**) across all samples. Two-way ANOVA was used for statistical significance testing (* p-value < 0.05) with p-value reported for the genotype effect (gRNA). Error bars are shown as SEM. (**K**) Boxplot showing the normalized molar percentages of the CE-to-free cholesterol (FC) ratio across all samples. (**L**) Boxplot showing the normalized molar percentages of the CE-to- FC ratio across all samples from both hiPSC lines grouped by genotype (CTRL = FA10 CTRL and IMR90 NT, KO/KD = FA10 TREM2 KO and IMR90 TREM2 KD). Statistical analysis for boxplots was performed using Mann Whitney test for non-parametric data (*** p-value < 0.001). Data was obtained from 1 experiment.
